# Impaired parvalbumin interneurons in the retrosplenial cortex as the cause of sex-dependent vulnerability in Alzheimer’s disease

**DOI:** 10.1101/2023.06.22.546142

**Authors:** Dylan J. Terstege, Yi Ren, Bo Young Ahn, Heewon Seo, Alzheimer’s Disease Neuroimaging Initiative, Liisa A. M. Galea, Derya Sargin, Jonathan R. Epp

## Abstract

Alzheimer’s disease is a debilitating neurodegenerative disorder with no cure and few treatment options. In early stages of Alzheimer’s disease, impaired metabolism, and functional connectivity of the retrosplenial cortex strongly predict future cognitive impairments. Therefore, understanding Alzheimer’s disease-related deficits in the retrosplenial cortex is critical for understanding the origins of cognitive impairment and identifying early treatment targets. Using the 5xFAD mouse model, we discovered early, sex-dependent alterations in parvalbumin-interneuron transcriptomic profiles. This corresponded with impaired parvalbumin-interneuron activity, which was sufficient to induce cognitive impairments and dysregulate retrosplenial functional connectivity. In fMRI scans from patients with mild cognitive impairment and Alzheimer’s disease, we observed a similar sex-dependent dysregulation of retrosplenial cortex functional connectivity and, in post-mortem tissue from subjects with Alzheimer’s disease, a loss of parvalbumin interneurons. Reversal of cognitive deficits by stimulation of parvalbumin interneurons in the retrosplenial cortex suggests that this may serve as a promising novel therapeutic strategy.

**One Sentence Summary:** Altered function, connectivity, and survival of parvalbumin expressing neurons in the retrosplenial cortex in Alzheimer’s Disease.

## INTRODUCTION

Alzheimer’s disease (AD) is associated with hallmark brain pathology including progressive appearance of amyloid plaques and neurofibrillary tangles. Even prior to development of these pathologies, numerous changes can be identified in brain structure and function. One of the earliest functional changes in individuals with mild cognitive impairment (MCI), who later develop AD, is hypometabolism in the posterior cingulate and, especially, the retrosplenial cortex (RSC)(*1–3*). This can occur years prior to the onset of AD. The degree of metabolic impairment in the RSC has been shown to predict the conversion from MCI to AD(*4*). Not all individuals with MCI will transition to AD. Determining those individuals who are most likely to convert is an important strategy that allows for early intervention. While numerous approaches such as cognitive testing and PET imaging of amyloid and tau have been used to identify at risk individuals, we have previously shown that RSC hypometabolism may be a superior risk factor. Recent work has indicated that RSC hypometabolism can predict MCI to AD conversion even in individuals that do not yet exhibit tau or amyloid pathology(*5*). This underlies both the importance of the RSC in disease progression and, it’s utility as a biomarker for AD progression and cognitive decline. To harness this utility, it is necessary to understand how changes that occur in the RSC lead to cognitive impairments. Little is known about the implications of the structural and functional changes that occur in the RSC. Previous work has demonstrated, that the RSC is a highly interconnected region critical for numerous forms of cognition(*6–8*). This, in conjunction with the early impairments in the RSC, warrant further investigation.

Here, we performed an in-depth analysis of the RSC in a transgenic mouse model of AD. We demonstrate that the activity and transcriptomic profile of the RSC is disrupted in 5xFAD mice. Our results point to impaired inhibition due to dysregulation and selective vulnerability of fast-spiking parvalbumin interneurons (PV-INs). Consequently, there is a disruption of global functional connectivity and cognitive performance. In healthy mice, these same pathological and cognitive impairments can be induced by selective inhibition of RSC PV-INs. Chemogenetic stimulation of the remaining PV-INs in 5xFAD mice is sufficient to enhance memory performance. Importantly, we found that the PV-IN impairments occur to a greater extent and earlier in females compared to males. This may explain the earlier onset of cognitive impairments in females with AD(*9*). To confirm these findings in AD patients, we used the ADNI neuroimaging database and found increased anticorrelated RSC functional connectivity in MCI and AD occurring earlier in females than males. In post-mortem AD tissue we also observed loss of PV-INs in the human RSC. Thus, our experiments demonstrate that RSC PV-INs are critical for maintaining healthy cognitive function and are selectively vulnerable in AD, especially in females. Targeting these neurons provides a therapeutic approach to prevent cognitive impairment.

## RESULTS

### Early Sex-Dependent Disruption of RSC Activity Despite Comparable Amyloid Pathology

Hypometabolic function in the RSC is observed early in clinical MCI/AD pathogenesis(*4*). To explore its implications, we confirmed disrupted metabolic activity in the RSC of 6-month-old 5xFAD mice (Fig. 1A, 1B, S1A). Cytochrome C oxidase labeling revealed reduced metabolic activity in female, but not male, 5xFAD mice (Fig. 1B). This sex-dependent difference was also noted with the neuronal glucose transporter GLUT3 (Fig. S1A). These data indicate that middle-aged female but not male 5xFAD mice, exhibit early RSC hypometabolism. However, we found comparable amyloid pathology in the RSC of both female and male 5xFAD mice (Fig. S1C)(*10*).Given the link between hypometabolism and neuronal hyperexcitability, supported by fMRI studies in AD patients(*11, 12*), we predicted increased RSC excitability in female 5xFAD mice. c-Fos immunohistochemistry following contextual fear conditioning revealed increased RSC neuronal activity in females but not in males (Fig. 1C, Fig. S1D).

**Fig. 1:**
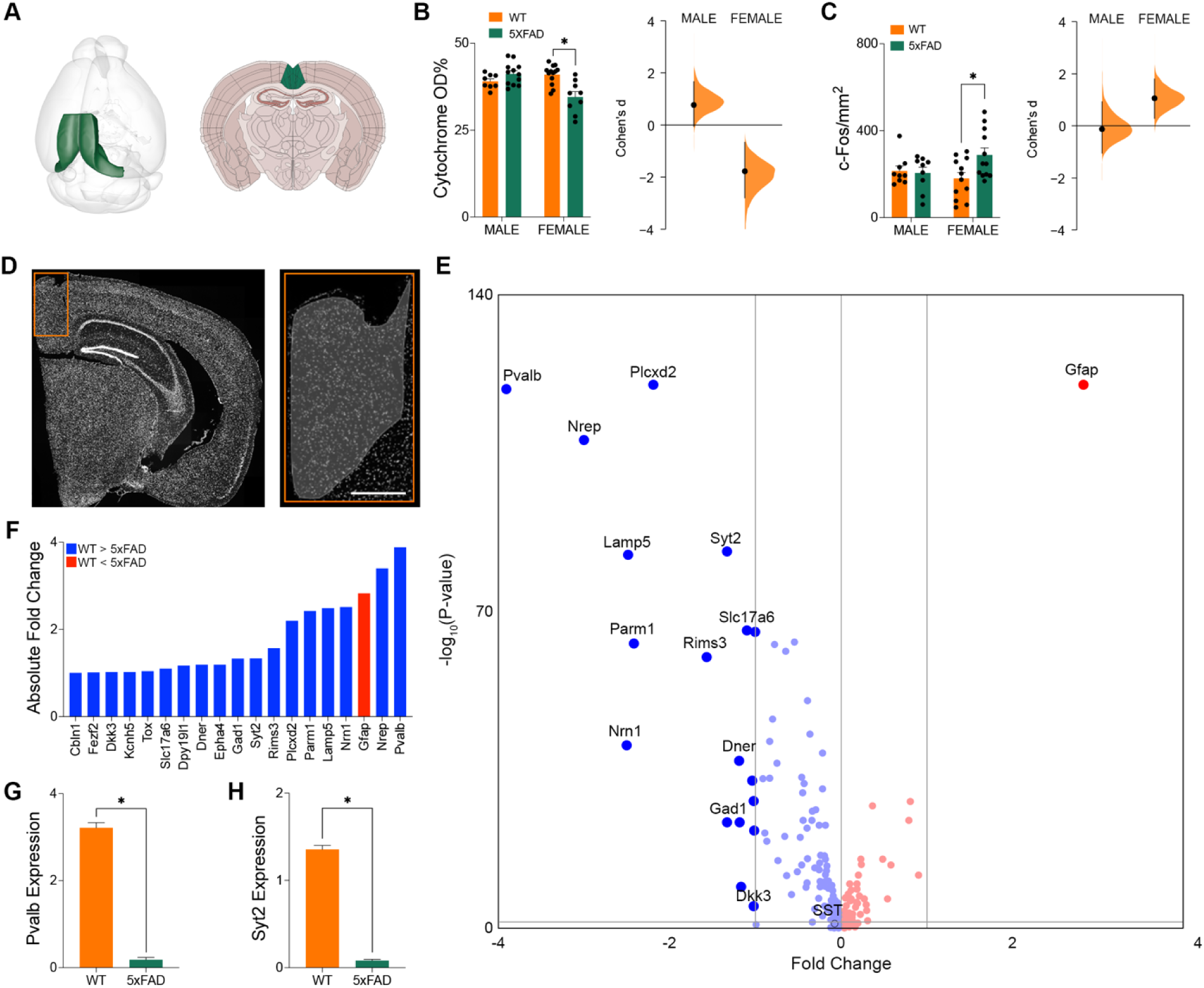
Hypometabolism, hyperexcitability, and downregulated *Pvalb* expression in the RSC of female 5xFAD mice. (**A**) RSC in the adult mouse brain. (**B**) Optical density of cytochrome C oxidase staining in the RSC. *P*=0.0003, Two-way ANOVA with Tukey’s multiple comparison test. Cohen’s D effect size between WT and 5xFAD for males (0.768) and females (−1.75). (**C**) Density of c-Fos+ cells in the RSC. *P*=0.0463, Two-way ANOVA with Tukey’s multiple comparison test. Cohen’s D effect size between WT and 5xFAD for males (−0.126) and females (1.06). (**D**) Nuclei labelled for single nucleus RNA sequencing across the RSC. Shaded region denotes area of interest. Scale bars represent 250μm. (**E**) Volcano plot of differentially expressed genes across the RSC of 5xFAD and WT mice. Dark blue, significantly downregulated in 5xFAD; dark red, significantly upregulated in 5xFAD; light colors, non-significant differential expression. (**F**) Absolute fold changes for all differentially expressed genes between WT and 5xFAD mice across the RSC. Normalized (**G**) *Pvalb* and (**H**) *Syt2* expression in the RSC of 5xFAD mice. *Pvalb*: adjusted *P*<0.0001, FDR-corrected (95%) Wilcoxon rank-sum test. *Syt2*: adjusted *P*<0.0001, FDR-corrected (95%) Wilcoxon rank-sum test. Data represent mean ± SEM. See also Figure S1. All statistical comparisons have been provided in Table S3.

### Downregulation of PV-IN Related Genes in the RSC of Female 5xFAD Mice

Following the confirmation of pathological changes in the RSC of 5xFAD mice, we investigated regional abnormalities using spatial transcriptomics (Fig. 1D). In comparing gene expression between female 6-month-old WT and 5xFAD mice, we identified 19 genes with significant differential expression (a nominal *p* <0.05 and log2 fold-change greater than ±1; Fig. 1E, 1F). Key genes involved in synaptic (*Kcnh5, Slc17a9, Rims3*) and metabolic (*Plcxd2, Nrn1, Nrep*) activities were downregulated, potentially explaining early changes in RSC hyperactivity and hypometabolism. The most significant change was the downregulation of *Pvalb*, which encodes the calcium-binding protein parvalbumin; a marker for a major class of fast spiking inhibitory interneurons (PV-INs). Interestingly, we also noted downregulation of several other genes that are normally highly enriched in PV-INs (*Syt2, Lamp5*) or in inhibitory interneurons in general (*Gad1*; Fig. 1G, 1H). Genes typically expressed in other inhibitory interneuron classes (e.g., calbindin, somatostatin, vimentin) were not significantly altered in 5xFAD mice. Together these data suggest a potential impairment in the RSC PV-IN population. Impairments in PV-INs, which could arise from disruptions of regional metabolic or activity, may have significant consequences for RSC function and cognitive performance.

### Early Sex-Dependent Disruptions Specific to RSC PV-INs

Given these transcriptional changes, we examined whether a corresponding loss of PV-INs occurred in the RSC (Fig. 2). Immunohistochemical labeling for parvalbumin showed a significant reduction of PV-INs in 6-month-old female 5xFAD mice, but no difference in males (Fig. 2A, 2B). Prior studies have reported the loss of inhibitory interneurons, including PV-INs, in several brain regions of AD patients and transgenic AD mice(*13–15*). We also investigated whether this loss extends to other brain regions with early AD pathology. We quantified PV-INs in the entorhinal cortex, subiculum, and CA1 of the hippocampus in 5xFAD mice(*16*). At 6 months, PV-INs were unaffected in both sexes across these other regions, although females generally had fewer PV-INs than males (Fig. S2A). By 12 months, PV-IN loss was evident in both sexes across multiple regions, but females showed a premature loss in the RSC compared to males as well as in other brain areas (Fig. S2B).

**Fig. 2:**
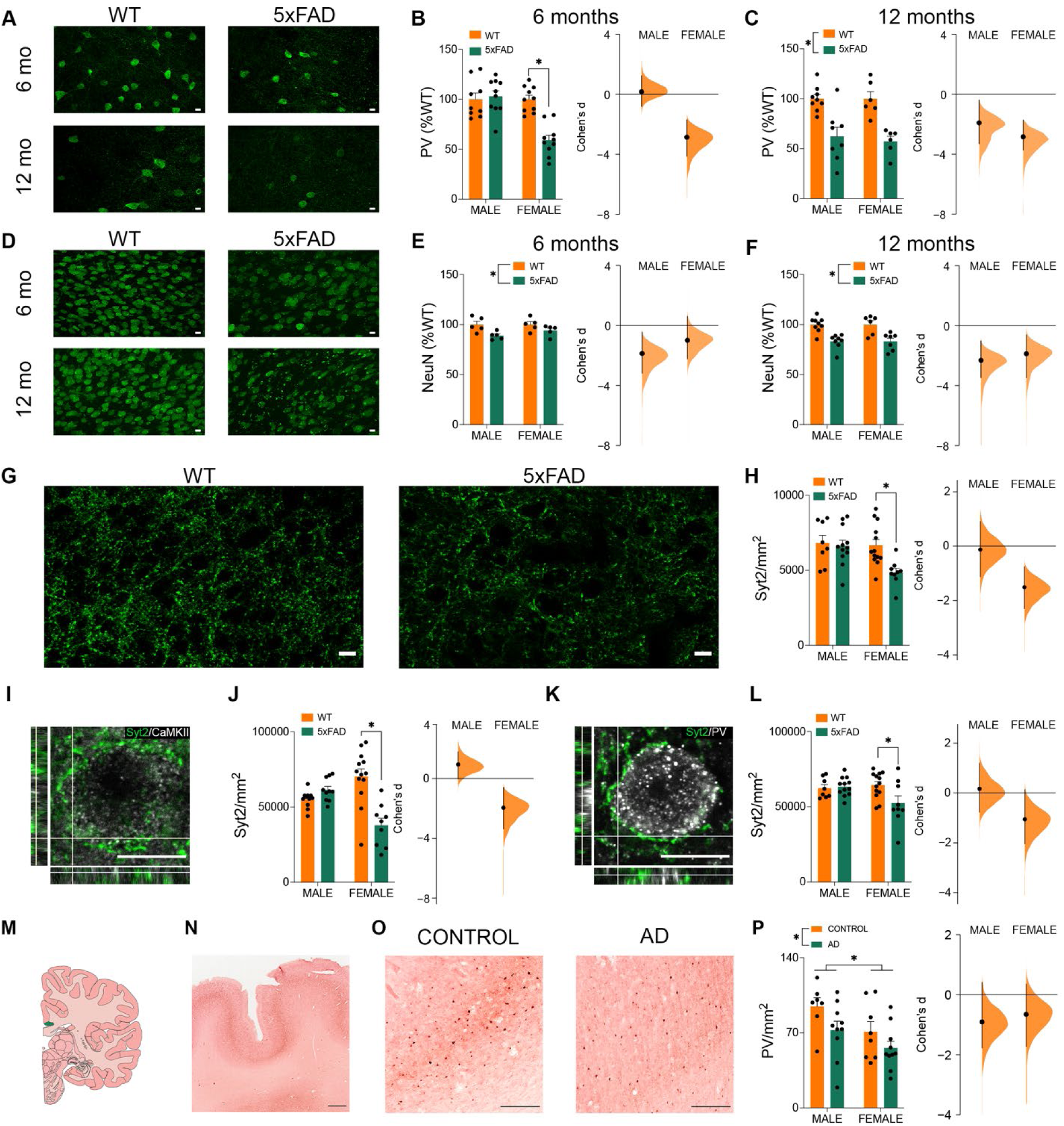
Immunohistochemical changes in the expression of PV-INs in the RSC. (**A**) PV staining in the RSC of 6- and 12-month-old WT and 5xFAD mice. Scale bars represent 10μm. RSC PV-IN density expressed as a percentage of the mean density in WT mice. (**B**) 6-months-old; *P*=0.0019, Two-way ANOVA, Tukey’s multiple comparison test. Effect size between WT and 5xFAD for males (Cohen’s D=0.166) and females (Cohen’s D=-2.86). (**C**) 12-months old; *P*<0.0001, Two-way ANOVA. Cohen’s D effect size between WT and 5xFAD for males (−1.91) and females (−2.84). (**D**) NeuN staining in the RSC of 6- and 12-month-old WT and 5xFAD mice. Scale bars represent 10μm. RSC NeuN+ cell density expressed as a percentage of the mean in WT mice. (**E**) 6-months-old; *P*=0.005, Two-way ANOVA. Effect size between WT and 5xFAD for males (Cohen’s D=-1.87) and females (Cohen’s D=-0.994). (**F**) 12-months-old; *P*<0.0001, Two-way ANOVA. Cohen’s D effect size between WT and 5xFAD for males (−2.31) and females (−1.88). (**G**) Synaptotagmin-2 (Syt2) staining in female WT and 5xFAD mice. Scale bars represent 10μm. (**H**) Syt2 density across the RSC. *P*=0.0111, Two-way ANOVA, Tukey’s multiple comparison test. Cohen’s D effect size between WT and 5xFAD for males (−0.134) and females (−1.51). (**I**) Syt2 (green) and CaMKII (greyscale) staining. Scale bars represent 10μm. (**J**) Syt2 density on CaMKII+ cells in the RSC. *P*<0.0001, Two-way ANOVA, Tukey’s multiple comparison test. Cohen’s D effect size between WT and 5xFAD for males (0.962) and females (−1.95). (**K**) Syt2 (green) and PV (greyscale) staining. Scale bars represent 10μm. (**L**) Syt2 density on PV-INs in the RSC. *P*=0.0254, Two-way ANOVA, Tukey’s multiple comparison test. Cohen’s D effect size between WT and 5xFAD for males (0.167) and females (−1.06). (**M**) Location of the human RSC (BA 29 & 30). PV staining in the human RSC. (**N**) Overview and (**O**) zoomed. Scale bar represents 250μm. (**P**) Density of PV-INs across the RSC. Main effects of sex (*P=*0.0068) and AD (*P=*0.0106). Two-way ANOVA. Cohen’s D effect size between control and AD groups for males (−0.906) and females (−0.654). Data represent mean ± SEM. See also Figure S2. All statistical comparisons have been provided in Table S3.

To distinguish whether loss of PV-INs in the RSC is due to generalized increase in neuronal death or is specific to PV-INs, we quantified the pan-neuronal marker NeuN (Fig. 2D – 2F). The approximate 40% loss of PV-INs in female 5xFAD mice contrasted with only a 6% decrease in overall RSC neurons, suggesting specific vulnerability of PV-INs in the female RSC. At 12 months, both males and females showed similar percentages of PV-IN loss (38% and 43%, respectively), which remained disproportionate compared to general neuronal loss.

### Reduced PV-IN Connectivity in the RSC of Female 5xFAD Mice

We hypothesized that a reduction in PV-INs would correspond to fewer synaptic contacts from PV-INs. This hypothesis was supported by our Xenium transcriptomics data showing downregulation of *Syt2*, a presynaptic marker selective for PV-INs (Fig. 1H).

Immunohistochemistry confirmed a significant decrease in the overall density of Syt2 protein labeling in female but not male 5xFAD mice (Fig. 2G, 2H). Specifically, Syt2 presynaptic terminals contacting CaMKII neurons and PV-IN cell bodies were reduced in females (Fig. 2I – 2L), with the magnitudes of these effect sizes indicating a preferential loss of inhibition in excitatory neurons, consistent with increased RSC excitability in female 5xFAD mice (Fig. 1D, S1).

### Validation of PV-IN Deficits as a Factor in AD

Our findings demonstrate sex-dependent vulnerability of PV-INs in the 5xFAD model, contributing to cognitive impairments. We confirmed that this result is not unique to 5xFAD mice by investigating the TgCRND8 model, where we observed a similar early PV-IN loss in female but not male TG+ mice (Fig. S2D, S2E)(*17*).

To establish whether similar PV-IN deficits occur in human Alzheimer’s Disease, we analyzed post-mortem RSC tissue from AD patients and age-matched controls. Our findings revealed a significant reduction in PV-INs in the RSC of AD patients compared to controls. Results from this analysis which used tissue from patients in advanced stages of AD aligns with our analysis of 5xFAD tissue from 12-month-old mice. At this stage both male and female patients (and mice) show a loss of PV-INs in the RSC. Intriguingly, in both humans and mice, female controls had fewer PV-INs compared to males. As a result, female AD patients (and transgenic mice) have significantly fewer PV-INs remaining in the RSC than male AD patients (Fig. 2M – 2P).

### Spatial Transcriptomics Demonstrates Enhanced Vulnerability of Female PV-INs

To investigate mechanisms underlying early and increased PV-IN vulnerability in females, we conducted cell-type-specific spatial transcriptomics using the Nanostring GeoMX platform. Both NeuN and PV were used as morphology markers to profile these neuron populations (PV+ neurons versus PV-neurons) (Fig. 3). This analysis revealed significant differential gene expression between 6-month-old WT and 5xFAD mice, particularly in female PV-INs, with a greater number of DEGs compared to males indicating that the greatest dysregulation may be occurring in female PV-INs (Fig. 3B). PV-INs in female 5xFAD mice exhibited downregulation of *Pvalb*, *Syt2*, *Gad1*, and *Arx*, all of which are critical to PV-IN function (Fig. 3C). These changes did not occur in PV-INs of male 5xFAD mice. In contrast, we also looked at a similar array of characteristic principal neuron genes including *NeuN* (neuron specific transcription factor), *Syp* (neuronal synaptic protein), *Vglut* (glutamate synthesis) and *Emx1* (regulator of pyramidal cell development) and found that none of these neuronal markers were differentially expressed in either male or female 5xFAD neurons (Fig. 3C). These results demonstrate a greater degree of vulnerability of female PV-INs at this earlier stage compared to the overall neuronal population (Fig. S3) and compared to male PV-INs.

**Fig. 3:**
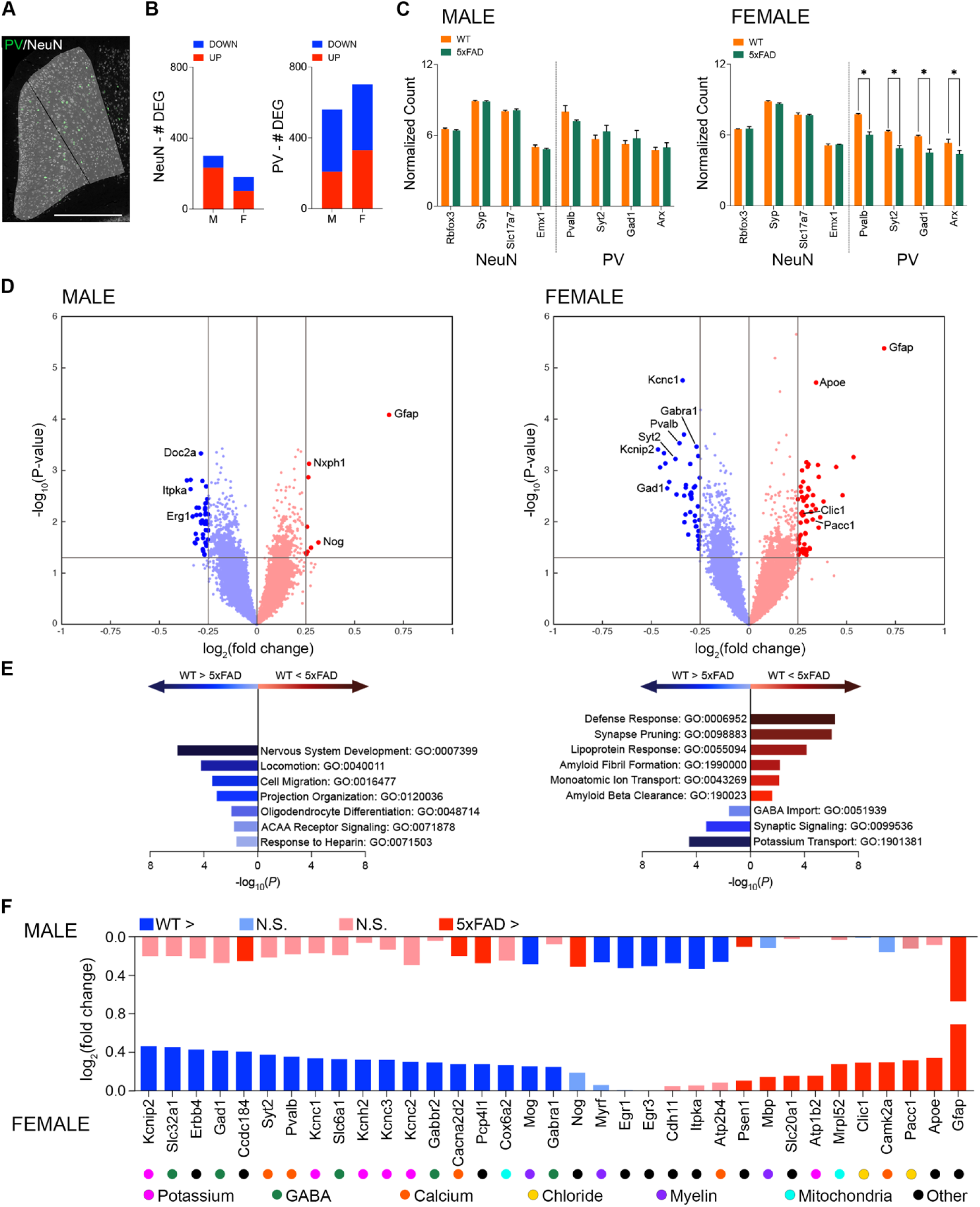
Altered transcriptomic profile in PV-INs of female 5xFAD mice. (**A**) PV (green) and NeuN (greyscale) staining in the RSC for GeoMX DSP analyses. Shaded regions denote areas of interest. Scale bars represent 500μm. (**B**) Number of differentially expressed genes across the RSC of male (M) and female (F) 5xFAD and WT mice. Nuclei isolated using NeuN staining (left) and PV staining (right). (**C**) Normalized counts for selected genes expressed by neurons and PV-INs from nuclei isolated using the NeuN and PV staining, respectively. *Pvalb* adjusted *P*=0.0003, *Syt2* adjusted *P*=0.0006, *Gad1* adjusted *P*=0.0022; *Arx* adjusted *P*=0.0178, FDR-corrected (95%) Wilcoxon rank-sum test. (**D**) Volcano plots of differentially expressed genes (DEG) across the RSC of 5xFAD and WT mice. Dark blue, significantly downregulated in 5xFAD; dark red, significantly upregulated in 5xFAD; light colors, non-significant differential expression. FDR-corrected (95%) Wilcoxon rank-sum test. (**E**) Top GO terms (by fold enrichment) identified by examining the top and bottom 30 DEG in PV-INs between male (left) and female (right) 5xFAD and WT mice. Blue, significantly downregulated gene clusters in 5xFAD relative to WT; red, significantly upregulated gene clusters in 5xFAD relative to WT. (**F**) Log_2_ fold changes observed between 5xFAD and WT mice in several selected genes in males (top) and females (bottom). Dark blue, significantly downregulated in 5xFAD; dark red, significantly upregulated in 5xFAD; light colors, non-significant differential expression. Genes involved in common processes have been grouped together according to the circular icons underneath of the gene names. Data represent mean ± SEM. See also Figure S3. All statistical comparisons have been provided in Table S3.

Gene ontology analysis showed that female 5xFAD PV-INs had altered expression of genes involved in cellular defense, synapse pruning, and amyloid-beta clearance, as well as decreased expression in GABA signaling and potassium transport (Fig. 3E; highlighted at the level of individual genes in Fig. 3F). These changes predict impaired physiological function in female PV-INs, contributing to RSC dysfunction.

### Reduced Inhibitory Control from PV-INs in the RSC of Female 5xFAD Mice

To determine the physiological alterations, we performed whole-cell patch-clamp electrophysiology in the RSC of 5xFAD mice. This revealed increased excitability in RSC pyramidal neurons in 6-month-old female 5xFAD mice compared to WTs, with females showing a much larger increase (Fig. 4A – 4F). While there was a small but significant increase in pyramidal cell excitability in male 5xFAD mice, the effect was consistently larger at all current steps in female 5xFAD mice compared to WT. With the exception of decreased capacitance in female 5xFAD mice there were no differences in any of the measured passive membrane properties of pyramidal cells in female or male 5xFAD mice (Fig. S4A – S4D). We observed a significant decrease in inhibitory postsynaptic current (sIPSC) amplitude in female but not male 5xFAD mice suggesting disrupted inhibition of pyramidal neurons in the female 5xFAD RSC (Fig. 4G – 4N). PV-INs play an important role in regulating pyramidal cell excitability(*18–20*). Therefore, to determine whether RSC hyperexcitability could relate to impaired PV-IN activity, we examined the electrophysiological properties of the RSC PV-INs. In male 5xFAD mice, we observed no significant differences in PV-IN excitability (Fig. 4O – 4R). However, in female 5xFAD mice, we found significant impairments in the activity of PV-INs (Fig. 4S – 4V).

**Fig. 4:**
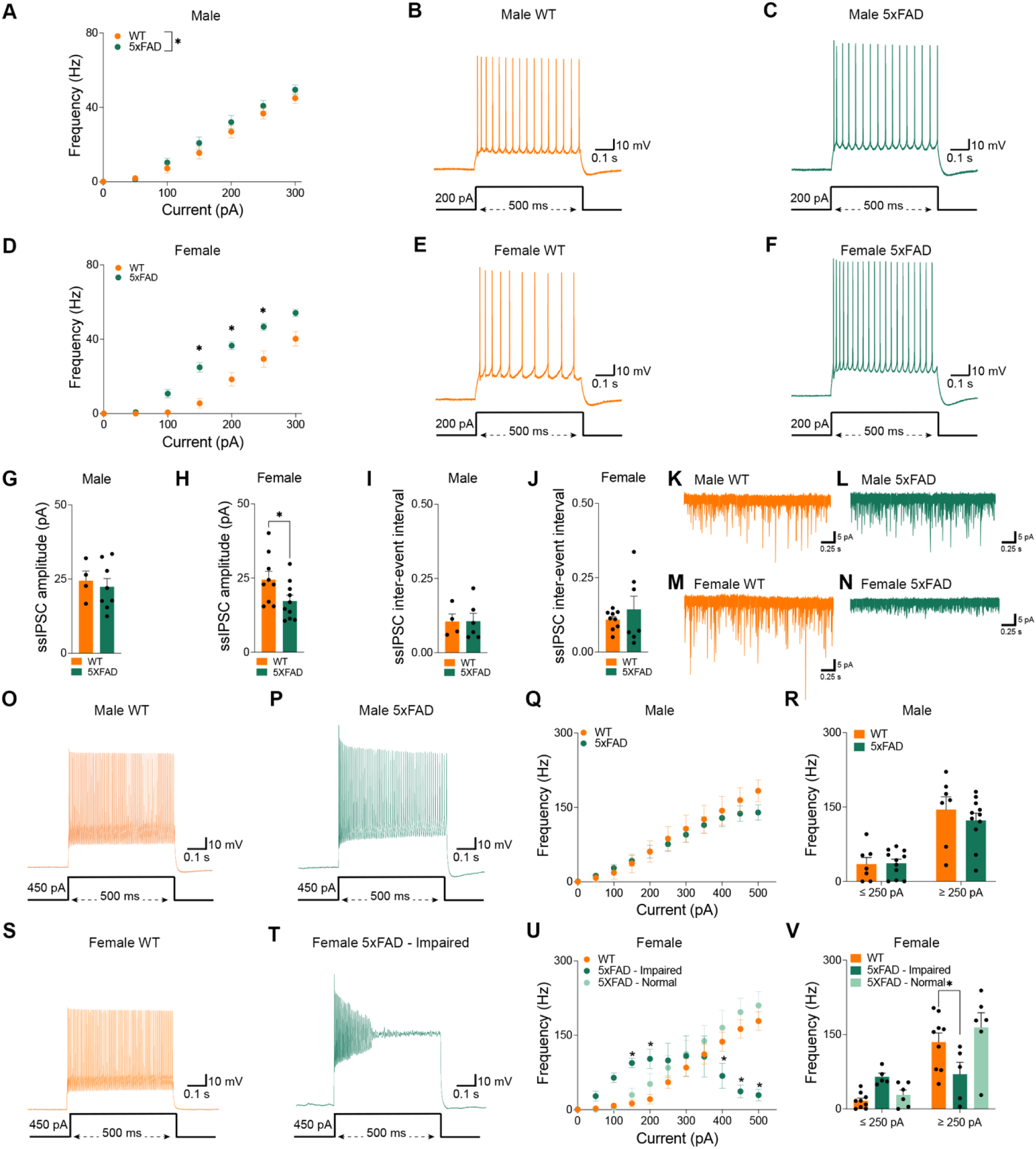
Hyperexcitable pyramidal neurons and impaired PV-IN activity in the RSC of female 5xFAD mice. (**A**) Firing frequency of pyramidal cells in male mice. *P=*0.0273, Two-way repeated measures ANOVA. Current-clamp traces of representative pyramidal cells from a male (**B**) WT and a (**C**) 5xFAD mouse in response to 200pA depolarizing current input. (**D**) Firing frequency of pyramidal cells in female mice. *P<*0.0001, Two-way repeated measures ANOVA. Current-clamp traces of representative pyramidal cells from a female (**E**) WT and a (**F**) 5xFAD mouse in response to 200 pA depolarizing current input. sIPSC amplitude in RSC pyramidal neurons in (**G**) male and (**H**) female WT and 5xFAD mice. *P*=0.0486, Two-sample t test. sIPSC inter-event interval in RSC pyramidal neurons in (**I**) male and (J) female WT and 5xFAD mice. Example sIPSC traces for (**K**) male WT, (**L**) male 5xFAD, (**M**) female WT, and (**N**) female 5xFAD mice. Current-clamp traces from PV-INs in male (**O**) WT and (**P**) 5xFAD mice in response to a 450pA depolarizing current input. (**Q**) Firing frequency of PV-INs in male WT and 5xFAD mice in response to depolarizing current injections. (**R**) Firing frequency of PV-INs in male mice in response to ≤250pA and ≥250pA current input. Current-clamp traces from (**S**) a PV-IN of a WT mouse and (**T**) an impaired PV-IN in a 5xFAD mouse in response to a 450pA depolarizing current input. (**U**) Firing frequency of PV-INs in response to depolarizing current injections. Dichotomous responses from two different PV-IN populations in 5xFAD females are visible (“5xFAD – Impaired” vs “5xFAD – Normal”). (**V**) Firing frequency of PV-INs in female mice in response to ≤250pA and ≥250pA current input. *P=*0.0252, Two-way repeated measures ANOVA, Dunnett’s multiple comparison test. Data represent mean ± SEM. See also Figure S4. All statistical comparisons have been provided in Table S3.

Interestingly, the pattern of impairment across PV-INs was not homogeneous. Instead, we observed two discrete populations based on their activity profiles across the I/O curve (Fig. 4U). One population of PV-INs (termed “impaired”) accounted for approximately half of the recorded neurons. These cells had elevated firing frequency at low depolarizing current inputs but then enter a state of depolarization block at larger current inputs (> 250 pA). In contrast, the other population exhibited normal firing frequencies at all depolarizing current inputs (Fig. 4V). We identified differences in passive membrane properties of PV-INs (input resistance, capacitance, rheobase current), but only in the impaired PV-IN population in 5xFAD females (Fig. S4E – S4L). There were no significant differences in the membrane characteristics of the normal PV-IN population of female 5xFAD mice, or PV-INs of male 5xFAD mice, compared with WTs. Although not all PV-INs are affected in 5xFAD female mice, it is evident that there is a broad impairment in the normal activity pattern of PV-INs, along with a greater reduction in the inhibitory control of pyramidal neurons in 5xFAD females, which does not occur in males.

### *In Vivo* RSC PV-IN Activity is Impaired in Female 5xFAD Mice

To determine *in vivo* RSC PV-IN population activity in 5xFAD mice, we performed calcium imaging with fiber photometry during a forced alternation variant of the Y-maze (Fig. 5A – 5E). To record specifically from PV-INs, we used an AAV virus with the PV-specific promoter s5e2 to express GCaMP6f in the RSC. Mice were first habituated to two arms of the Y-maze. During the test trial, mice were allowed to explore all three arms. We found a significant sex by genotype interaction with a decrease in time spent in the novel arm in female (but not male) 5xFAD mice indicating impaired recognition memory (Fig. 5D). In WT mice we observed an increase in PV-IN population calcium activity in the RSC during the transition between familiar and novel arms. However, the magnitude of this signal was significantly reduced in 5xFAD mice (Fig. 5C, 5E), and the effect size was larger in females than it was in males. Overall, these data show impaired PV-IN activity in the RSC during behavioral performance, which is a particularly robust finding in female 5xFAD mice.

**Fig. 5:**
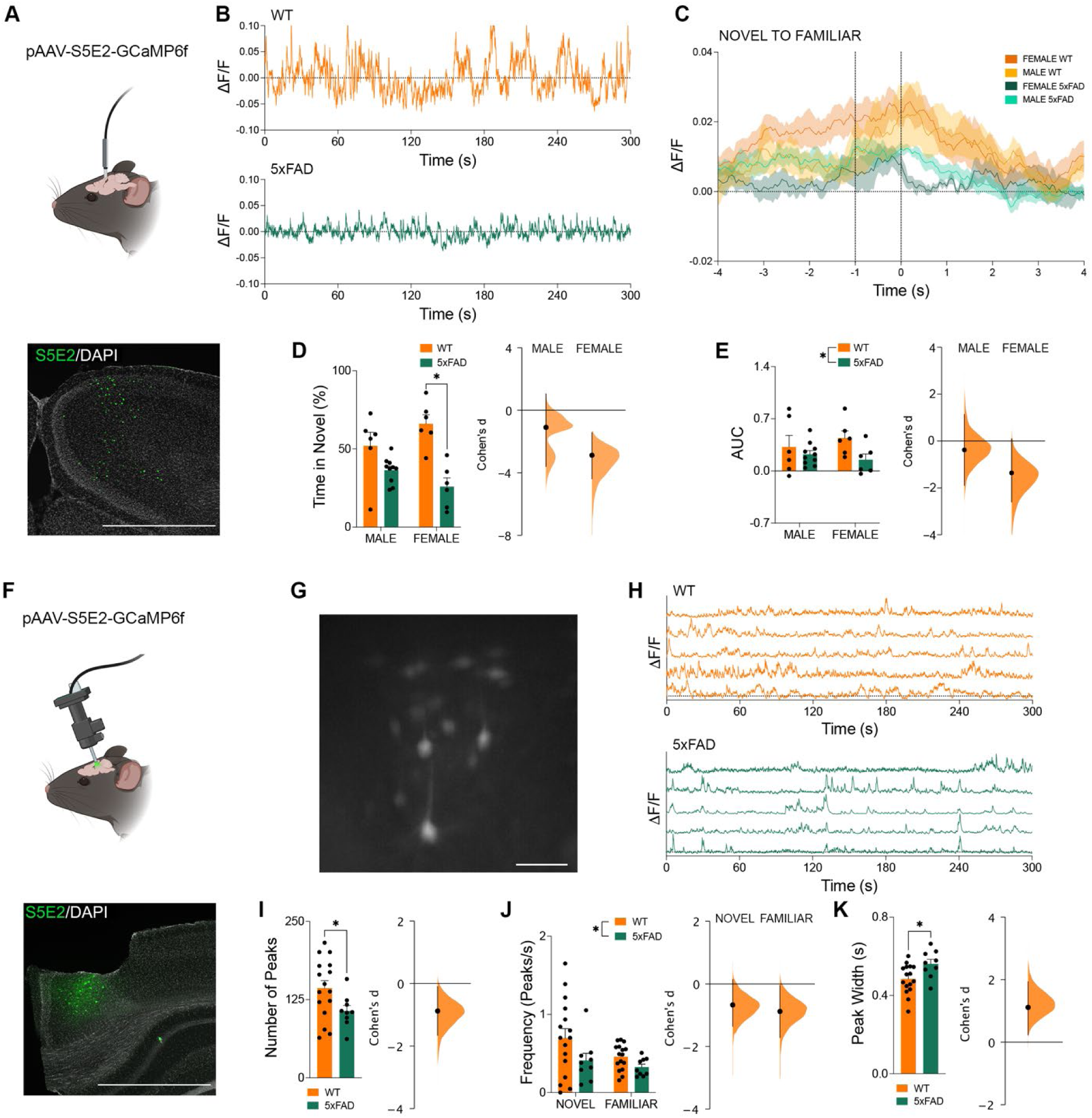
Impaired activity of RSC PV-INs *in vivo* during working memory performance. (**A**) pAAV-s5e2-GCaMP6f injection, optic ferrule placement, and virus expression (green). Scale bar represents 1mm. (**B**) ΔF/F traces showing the population activity of RSC PV-INs in freely behaving WT (top) and 5xFAD (bottom) mice. (**C**) ΔF/F trace during the 4s prior to and following familiar arm entry. (**D**) Percentage of time spent in the novel arm of the y maze. *P*=0.0003, Two-way ANOVA, Tukey’s multiple comparisons test. Cohen’s D effect size between WT and 5xFAD for males (−1.09) and females (−2.88). (**E**) Area under the curve (AUC) of the ΔF/F during the second preceding novel arm entry. *P*=0.047, Two-way ANOVA. Effect size between WT and 5xFAD for males (Cohen’s D=-0.381) and females (Cohen’s D=-1.36). (**F**) pAAV-s5e2-GCaMP6f injection, lens placement, and virus expression (green). Scale bar represents 1mm. (**G**) s5e2-GCaMP expression through the miniscope. Scale bar represents 50μm. (H) ΔF/F traces showing the activity of RSC PV-INs in freely behaving WT (top) and 5xFAD (bottom) mice. (**I**) Number of peaks in the ΔF/F signals of each cell during the 5-minute Y maze forced alternation task. *P*=0.0461, Two-sample t test. Cohen’s D effect size between WT and 5xFAD mice (−0.879). (**J**) Peak frequency in the ΔF/F signals of each cell in the novel and familiar arms of the Y maze forced alternation task. *P*=0.0301, Two-way ANOVA. Cohen’s D effect size between WT and 5xFAD in the novel arm (−0.672) and familiar arms (−0.882). (**K**) Peak width in the ΔF/F signals of each cell during the Y maze forced alternation task. *P*=0.0195, Two-sample t test. Cohen’s D effect size between WT and 5xFAD mice (1.11). Data represent mean ± SEM. All statistical comparisons have been provided in Table S3.

Fiber photometry reports difference in population activity but does not differentiate whether a change in activity is due to changes in individual neuron activity or changes in the number or synchronization of active neurons. Therefore, to determine whether individual PV-INs in the RSC show impaired activity *in vivo* we performed calcium imaging with head mounted miniscopes (Fig. 5F – 5K). Similar to what was observed with fiber photometry we found a significant decrease in the activity of individual RSC PV-INs in female 5xFAD mice during a forced alternation variant of the Y-maze. While performing this task, RSC PV-INs in female 5xFAD mice displayed reductions in both the overall number of peaks and peak frequency (Fig. 5I, 5J). We also observed a significant increase in the peak width in 5xFAD mice (Fig. 5K). Overall, these findings show impaired *in vivo* RSC PV-IN activity at both the single cell and population level.

### Sex-Dependent Disruption of RSC Functional Connectivity in 5xFAD Mice and in AD Patients

Our results to this point clearly indicate localized impairments in the activity of the RSC. However, the RSC also exhibits extensive structural and functional connectivity with numerous brain regions. The RSC is a critical hub in the brain’s default mode network, is highly involved in various types of cognitive processing and, functions as a bridge between cortical and subcortical brain regions(*21–23*). For these reasons, disrupted RSC PV-IN activity is likely to also disrupt communication with other brain regions and impair network activity broadly. Therefore, we investigated the functional connectivity of the RSC in 5xFAD mice following a contextual fear conditioning test (Fig. 6A – 6D, S5A – S5C). Using c-Fos immunolabelling we quantified regional activity in the RSC as a seed region and in an additional 90 brain regions. From this we cross correlated regional activities with the activity of the RSC as a measure of functional connectivity (Fig. S5B). A striking difference that emerged was a substantial increase in anti-correlated activity between the RSC and other brain regions. While the number of anti-correlated functional connections was low in control groups, it was substantially increased in female 5xFAD mice (Fig. 6B, 6C). Male 5xFAD mice did not show the same increase; and instead show a small decrease in anti-correlated connections. Additionally, we observed in female but not male 5xFAD mice, a decrease in local efficiency of the RSC (Fig. 6D; a graph theory metric indicative of the efficiency of information transfer between the RSC and its connected network(*24*)).

**Fig. 6:**
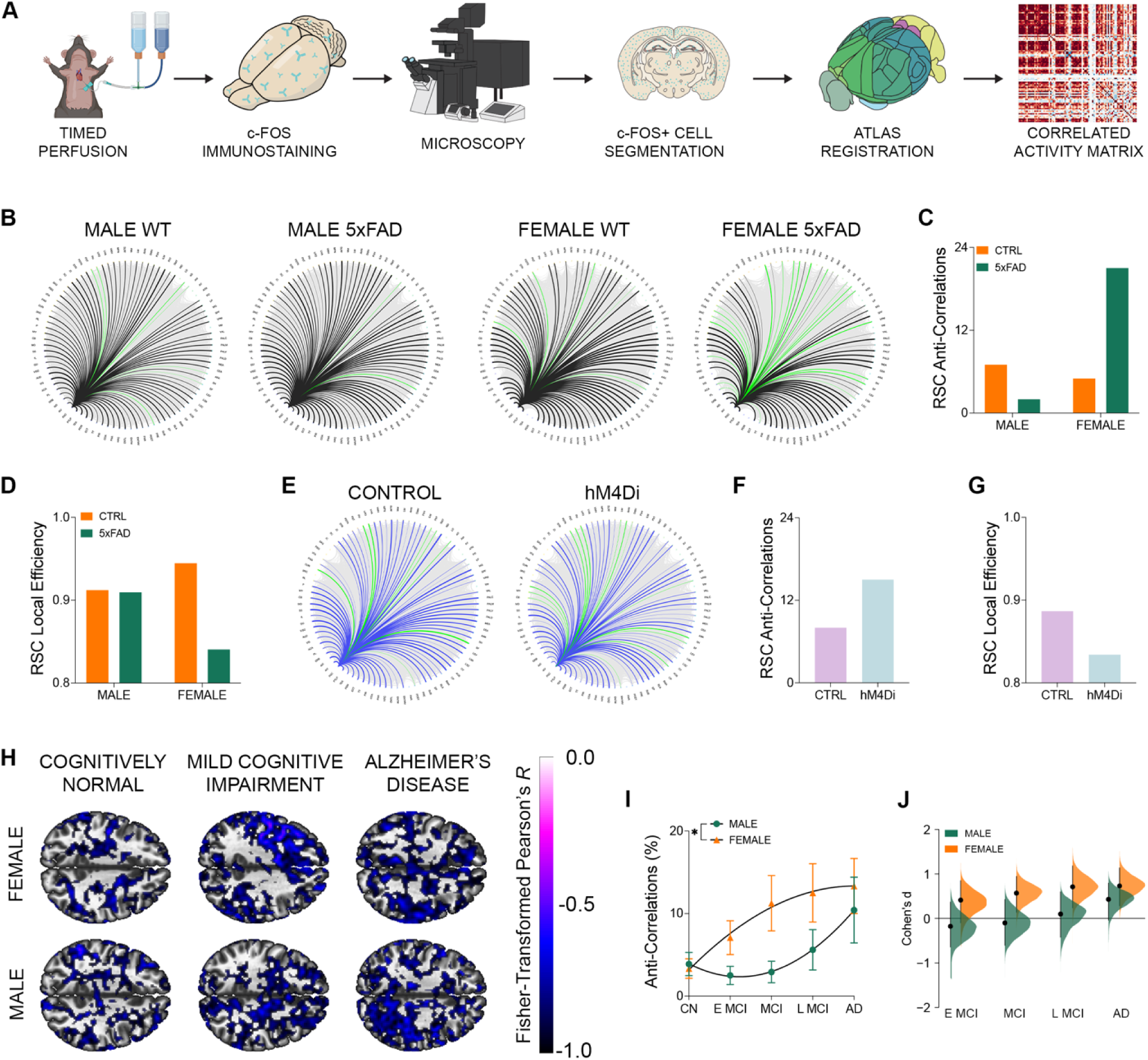
Pattern of altered RSC functional connectivity in female 5xFAD mice, healthy mice with dampened RSC PV-IN activity, and in clinical AD progression. (**A**) Schematic outlining the generation of functional connectivity networks from brain-wide c-Fos staining. (**B**) Circle plots of the RSC functional connectome in female WT, female 5xFAD, male WT, and male 5xFAD mice. Correlations with positive Pearson’s correlation coefficients are depicted in black. Correlations with negative correlation coefficients (anti-correlations) are depicted in green (**C**) Number of anti-correlations involving the RSC in male and female WT and 5xFAD mice. (**D**) Local efficiency of the RSC in male and female WT and 5xFAD mice. (**E**) Circle plots of the RSC functional connectome in control, and RSC s5e2-hM4Di injected mice. Correlations with positive Pearson’s correlation coefficients are depicted in blue. Anti-correlations are depicted in green. (**F**) Number of anti-correlations involving the RSC in control and RSC s5e2-hM4Di injected mice. (**G**) Local efficiency of the RSC in control and RSC s5e2-hM4Di injected mice. (**H**) Coactivation maps depicting resting state activity anti-correlated with the left RSC in a CN male, CN female, male with MCI, female with MCI, male with AD, and female with AD. (**I**) Nonlinear regression line showing the change in the density of anticorrelations involving the RSC within the DMN. *P=*0.0121, Sum of Squares F test. (**J**) Cohen’s D effect size difference in the density of anticorrelations within the DMN involving the RSC, in E MCI (males: −0.17; females: 0.409), MCI (males: −0.0999; females: 0.574), L MCI (males: 0.0941; females: 0.662), and AD (males: 0.429; females: 0.73) relative to sex-matched CN participants. Data represent mean ± SEM. Also see Figure S5. All statistical comparisons have been provided in Table S3.

Next, to establish whether similar disruptions in RSC functional connectivity could be observed during the clinical AD pathogenesis, we analyzed resting state fMRI data from the ADNI database. Using these data, we were able to As a key component of the default mode network (DMN, a network of functionally connected regions which becomes coactive while an individual is at rest), it was expected that the activity of the RSC should have very few anti-correlations with other DMN regions in rsfMRI scans of cognitively normal individuals(*7, 22, 25*). Seed-based functional connectivity analyses of the rsfMRI data revealed sex- and disease stage-specific changes in the density of anti-correlations within the DMN involving the RSC (Fig. 6H). The onset of impaired functional connectivity, in the form of increased anti-correlated activity of the RSC, followed a more rapid trajectory – from cognitively normal, through various stage of MCI, and into AD – in female patients than in males (Fig. 6I, J, S5N).

The impaired network activity of the RSC in female 5xFAD mice and AD patients suggests a dysregulation of communication between the RSC and other brain regions that may underlie observed cognitive impairments(*26*). In fact, intact functional connectivity of the RSC has been suggested to be a key indicator of healthy cognitive aging(*6*). Anti-correlated functional connectivity is often overlooked but, has been speculated to relate to activity driven by inhibitory interneurons(*27, 28*). If so, changes in anti-correlated activity would be predicted to occur based on the loss and disruption of PV-IN activity observed here. To our knowledge, the relationship between local inhibition and anti-correlated functional connectivity has never been directly investigated.

### Chemogenetic Inhibition of PV-INs Recapitulates AD Impairments

As a test of this relationship, we performed targeted inhibition of PV-INs in the RSC of healthy WT mice. We infected PV-INs with an AAV expressing the inhibitory chemogenetic DREADD, hM4Di under control of the s5e2 promoter that is highly selective for PV-INs (Fig. S5D – S5F. We predicted that chemogenetic inhibition of PV-INs would impair learning and memory and reproduce the increased RSC anti-correlated functional connectivity. Mice expressing hM4Di specifically in PV-INs (Fig. S5F – S5G) and treated with the ligand DCZ, showed significantly weaker contextual fear memory compared to mice infected with a control virus (Fig. S5H – S5I). This was true regardless of sex with an equal effect size observed in female and male mice. The lack of a sex difference in this experiment is not unexpected given that we experimentally induced the underlying impairment in PV-IN activity in both male and female mice. We next perfused the mice 90 minutes following memory retrieval to examine functional connectivity (Fig. 6E). Mice expressing hM4Di and treated with DCZ exhibited decreased PV-IN and increased RSC activity (Fig. S5J – S5L). Importantly, hM4Di mice displayed increased RSC anti-correlated connectivity (Fig. 6F) as well as a decrease in the local efficiency of the RSC (Fig. 6G), similar to the deficits in female 5xFAD mice. This suggests that the occurrence of increased anti-correlated activity in 5XFAD mice is related to disrupted PV-IN function. Importantly, these data show that impaired PV-IN activity is sufficient to induce anti-correlated functional connectivity and memory impairment.

### Stimulation of RSC PV-INs to Reverse Cognitive Deficits in Female 5xFAD Mice

Given the deficits in PV-IN function and the corresponding overall RSC hyperexcitability, we explored the use of targeted PV-IN stimulation to rescue behavioral impairments (Fig. 7). We used an AAV expressing the excitatory DREADD hM3Dq under the s5e2 promoter to target PV-INs in the RSC of 6-month-old 5xFAD mice. Three weeks after surgery, we performed behavioral tests to assess whether stimulation of RSC PV-INs would improve cognitive behavior (Fig. 7A). We administered DCZ before performing forced alternation in a Y-maze and before the retention test of a contextual fear conditioning task. In male mice, we did not see an effect of RSC PV-IN stimulation on Y-maze behavior (Fig. 7C). However, in female 5xFAD mice, RSC PV-IN stimulation led to increased recognition of the novel arm in the Y-maze forced alternation test (Fig. 7D). Similarly, contextual fear memory was not impacted by RSC PV-IN stimulation in male mice; while in female 5xFAD mice, contextual memory performance was significantly improved, reflected by a decrease in the activity suppression ratio, thereby indicating that mice spent less time moving upon context reintroduction than they did prior to shock delivery during training (Fig. 7E, 7F). To verify the effects of DCZ on the activity of s5e2-AAV labeled PV-INs, we assessed the activity of these cells with c-Fos colocalization. Compared to saline treatment, DCZ increased the activity of s5e2-AAV labeled PV-INs (Fig. 7G, 7H). We also observed a corresponding decrease in overall RSC c-Fos expression, indicating that the DREADD/DCZ stimulation successfully reduced the hyperexcitability of the region as a whole (Fig. 7I, 7J).

**Fig. 7:**
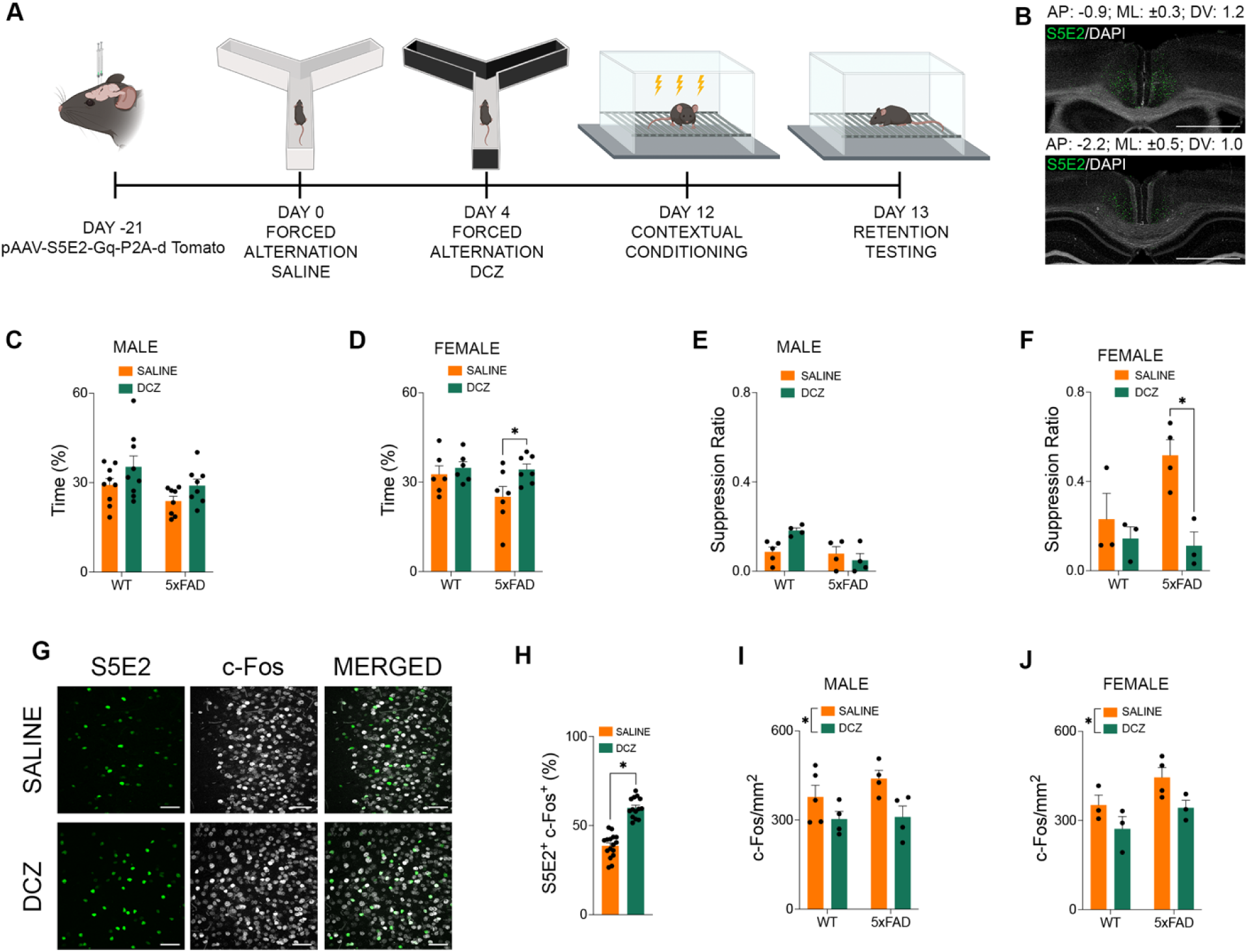
Stimulation of RSC PV-INs promotes the recovery of memory performance in 5xFAD mice. (**A**) Timeline of surgery and behavioral testing. (**B**) pAAV-s5e2-Gq-P2A-d Tomato expression (green) and DAPI (greyscale). Scale bars represent 1mm. (**C**) Percentage of time spent in the novel arm of the forced alternation Y-maze task by male mice with saline treatment and subsequent DCZ treatment. (**D**) Percentage of time spent in the novel arm of the forced alternation Y-maze task by female mice with saline treatment and subsequent DCZ treatment. *P*=0.0349, *a priori* Two-sample t test. (**E**) Activity suppression ratio following contextual conditioning by male mice treated with saline or DCZ. (**F**) Activity suppression ratio following contextual conditioning by female mice treated with saline or DCZ. *P*=0.0088, *a priori* Two-sample t test. (**G**) Photomicrographs showing s5e2 (green) and c-Fos (greyscale) staining in the RSC with saline and DCZ treatments. Scale bars represent 50μm. (**H**) Percentage of s5e2+ cells which were colocalized with c-Fos with saline and DCZ treatment. *P*<0.0001, Two-sample t test. (**I**) c-Fos expression density across the RSC in male WT and 5xFAD mice treated with saline of DCZ. *P=*0.0108, Two-way ANOVA. (**J**) c-Fos expression density across the RSC in female WT and 5xFAD mice treated with saline of DCZ. *P=*0.0260, Two-way ANOVA. Data represent mean ± SEM. Also see Figure S6. All statistical comparisons have been provided in Table S3.

## DISCUSSION

Despite early functional impairments in the RSC in AD, little work has characterized these pathological changes or their implications for cognitive performance. The RSC is critical for many cognitive functions due to its intrinsic processing and high degree of connectivity. Understanding the vulnerability of this region to AD-related pathology would be highly beneficial for early diagnostic and/or intervention strategies. Given the sex differences in the onset of cognitive impairments in AD, we reasoned that understanding these differences necessitates examining the earliest impacted brain regions. Our goal was to explore the functional deficits in female and male RSC using a mouse model of AD.

We confirmed that the hypometabolic/hyperactive phenotype observed in the human RSC is also evident in 5xFAD mice, particularly in females. We observed both increased excitability during memory retrieval and decreased metabolism. Intriguingly, these deficits were observed in female but not male 5xFAD mice. Using Xenium transcriptomics to investigate changes in the RSC, we identified a significant downregulation *of Pvalb* and PV-IN-related genes, which were among the largest changes in gene expression in 5xFAD mice. We followed-up on this observation with cell-type-specific transcriptomics guided by immunohistochemistry to compare differential gene expression within specific cellular populations in 5xFAD and WT mice, particularly between PV-IN populations. Here, we observed a greater number of differentially expressed genes in PV-INs, especially in females. Gene Ontology-based analysis indicated that the changes in gene expression in PV-INs could lead to impaired synaptic signaling, increased pruning, impaired GABAergic function, and alterations in neuronal excitability. These predictions are supported by our subsequent electrophysiology, calcium imaging, and histology analyses.

To investigate the accompanying physiological impairments in RSC, we performed whole-cell patch-clamp electrophysiology in female and male 5xFAD mice. We found increased excitability of RSC pyramidal cells, which was more prominent in female 5xFAD mice. Neuronal hyperexcitability is commonly observed in AD mice and is supported by imaging studies in patients with MCI and AD; however, these changes in excitability are not static in their presentation and show both regional susceptibility and temporal variability. In MCI patients, hyperactivity is observed in some brain regions, while AD is associated with hypoactivity in the same regions, suggesting a temporal progression from hyper- to hypo-excitability with disease progression. The underlying mechanisms of this transition are not completely understood. Our study suggests that the sex-dependent hyperexcitability in the RSC pyramidal neurons coincide with early impairments in PV-IN activity. This could lead to reduced inhibition within the RSC, which could explain the increased hyperexcitability of pyramidal neurons and the broadly increased c-Fos expression across the RSC in female 5xFAD mice. The emergence of two distinct PV-IN populations in female 5xFAD mice (one normal and one impaired) may represent different stages of degeneration or distinct subclasses of PV-INs that respond differentially during neurodegeneration.

PV-INs are crucial for regulating neuronal excitability. These interneurons synapse extensively onto pyramidal neurons and other inhibitory interneurons, including other PV-INs. Individual PV-INs synapse onto multiple pyramidal cells, each of which typically receives input from multiple PV-INs. The reduction in presynaptic PV-IN connectivity is significantly greater on CaMKII+ neurons than on other PV-INs. This differential loss of PV-IN synapses across various cell types, along with the reduction in inhibitory synaptic currents in RSC pyramidal neurons of female 5xFAD mice, further supports that the increased excitability of RSC pyramidal neurons is likely due to reduced inhibition from PV-INs.

In addition to our observation of disrupted PV-IN activity, we found significantly reduced PV-IN numbers in the RSC. This deficit in PV-IN expression was evident at a relatively early stage (6 months old) in female 5xFAD mice but was not present in age-matched males. By 12 months of age, however, PV-IN loss was observed in both female and male 5xFAD mice compared to controls. Notably, the loss of PV-INs in 6-month-old female 5xFAD mice was selective to the RSC and not observed in other brain regions with early AD pathology until mice reached 12 months of age. The reduction of PV-INs in other brain regions of 5xFAD mice by 12 months old is equivalent to the previously reported time course of PV loss in the hippocampus and coincides with impairments in gamma oscillatory behaviors that have been reported at this age(*29*). Importantly, the PV-IN population in the RSC shows greater vulnerability, significantly exceeding the slower rate of general neuronal loss observed in this area. Together, these findings indicate that PV-INs in the female RSC are an attractive target for early intervention due to their unique early vulnerability to AD.

Chemogenetic inhibition of PV-INs in healthy mice recapitulated the impairments observed in 5xFAD mice, including disruptions in RSC functional connectivity, regional excitability, and behavior. Chemogenetic stimulation of PV-INs in 5xFAD mice was able to rescue these impairments, further supporting the integral role of RSC PV-INs in AD-related disruptions. These findings provide proof of principle for targeted intervention to promote cognitive enhancement. Even at an advanced age of 12 months, corresponding to a relatively progressive stage in AD, targeting the RSC PV-INs was sufficient to enhance memory. This underscores the potential of improving the function of this specific cell type as a treatment for AD. However, the stimulation of the PV-IN population may be limited by the fact that disrupted PV activity in the RSC is accompanied by a reduction in the number and connectivity of PV-INs. Future studies should explore alternative approaches aimed at preventing the loss of PV-INs and preserving their connectivity, which could delay the onset of cognitive impairment, particularly in females where such deficits manifest earlier and more profoundly.

An important finding of this study is the sex-specificity of PV-IN dysfunction. The incidence of AD is two-fold greater, and the progression from MCI to AD is more rapid in women than men(*30, 31*). Despite these discrepancies, sex-stratified data from randomized clinical trials of AD treatments is scarce. Even in preclinical models, sex differences are often not assessed. Our results identify that, in both humans and mice, the baseline expression density of PV-INs is lower in in females. Furthermore, our results suggest an increased vulnerability of PV-INs to AD pathology at earlier disease stages. These data collectively demonstrate the potential of PV-INs as a target for intervention and the potential use of RSC neuroimaging as a clinical tool for tracking MCI/AD progression.

Bridging findings from preclinical animal models to clinical cases of AD can be challenging. However, this study shows commonalities in RSC impairments between 5xFAD mice and clinical AD across multiple features, including hypometabolism, PV-IN density, and impaired functional connectivity. These data also suggest different trajectories in AD progression, with impaired PV-IN density in 6-month-old 5xFAD female mice coinciding with impaired functional connectivity in women with MCI. This is further supported by the absence of sex differences in advanced stages of the disease, as evidenced by the similar PV-IN expression density, as well as the lack of sex differences in impaired functional connectivity in 12-month-old 5xFAD mice and patients clinically diagnosed with AD. These findings align with clinical differences in the rates of progression from MCI to AD between men and women. Taken together, these complementary preclinical and clinical data suggest that impaired PV-IN function in the RSC may be a mechanism contributing to sex differences in AD progression and prevalence. Importantly, even in 12-month-old 5xFAD exhibiting profound baseline cognitive impairments, stimulation of RSC PV-INs provides a mechanism for improving cognitive performance. Given the high degree of convergence between the impairments observed in the transgenic mouse model and the clinical AD population, our study provides a potential neurostimulation strategy for preserving cognitive ability in AD/MCI patients.

In conclusion, we have identified a novel impairment in PV-IN function in the RSC, a critically understudied brain region with respect to AD. Importantly, these findings mirror the earlier cognitive impairments observed in females. Our findings provide both a potential mechanism to explain why this sex difference occurs and a specific intervention target to enhance cognition. Protection of the vulnerable PV-IN cell population represents a key to slowing cognitive decline in MCI/AD patients.

## MATERIALS AND METHODS

### Study Design

We investigated the transcriptomic profile, cellular activity, and functional connectivity of a neuroanatomical region thought to be predictive of cognitive impairment in AD, the RSC. These investigations were performed using mouse models of AD, human tissue samples, and neuroimaging data from clinical populations. All experiments involving animal subjects were conducted in accordance with Canadian Council on Animal Care guidelines and with the approval of the University of Calgary Animal Care Committee. All experiments involving human tissue samples were conducted with the approval of the University of Calgary Conjoint Health Research Ethics Board (REB22-0776). All analyses were performed by experimenters blinded to experimental conditions.

### Subjects

#### Mice

Heterozygous 5xFAD mice were produced via *in vitro fertilization by the* University of Calgary Centre for Genome Engineering. Sperm was obtained from male 5xFAD mice (#034840-JAX) and female C57Bl/6J mice were used as oocyte donors (#000664-JAX). Male and female experimental offspring were aged to 6 or 12 months for use in most experiments. Experiments in which PV-IN activity was chemogenetically silenced were conducted using 8-week-old male and female C57Bl/6J mice. Further histological analyses were also conducted using male and female offspring from a TgCRND8 (MGI: 3589475) and C57Bl/6J cross, aged to 5 months. Mice were housed in standard cages with three to five mice per cage and free access to food and water. The room lighting was maintained on a 12 h/12 h light/dark cycle (8 am, lights on). All procedures were conducted during the light cycle phase.

#### Human tissue samples

Frozen-fixed RSC tissue from male and female patients with Alzheimer’s disease (Mean Age ± SD: Male 71.2 ± 3.3, Female 72.7 ± 2.6) and age-matched individuals without Alzheimer’s disease (Mean Age ± SD: Male 69.3 ± 2.8, Female 71.6 ± 3.0) was obtained from the Douglas-Bell Canada Brain Bank (Montreal, QC, CA). The known average duration of disease was 8.1 ± 3.4 years in Males AD patients and 7.4 ± 4.6 years in female AD patients. In collaboration with the Quebec Coroner’s Office and with informed consent from the next of kin, phenotypic information was obtained with standardized physiological autopsies. Control and Alzheimer’s disease were defined with the support of medical charts and Coroner records. Toxicological assessments, medication prescription, and cognitive assessments were also obtained.

#### Human neuroimaging datasets

Data used in the preparation of this manuscript were obtained from the Alzheimer’s Disease Neuroimaging Initiative (ADNI) database (adni.loni.usc.edu). The ADNI was launched in 2003 as a public-private partnership led by Principal Investigator Michael W. Weiner, MD. The primary goal of ADNI has been to test whether serial MRI, PET, other biological markers, and clinical and neuropsychological assessment can be combined to measure the progression of MCI and early AD. For up-to-date information, see http://www.adni-info.org.

Neuroimaging data from 114 participants was analyzed from the ADNI 1 and 2 databases. This includes 59 cognitively normal individuals (CN; *n*=29 males Mean Age ± SD: 73.3 ± 8.6, *n*=30 females Mean Age ± SD: 73.1 ± 8.7), 175 individuals experiencing mild cognitive impairment, and 60 individuals diagnosed with Alzheimer’s Disease (AD; *n*=30 males Mean Age ± SD: 73.0 ± 7.7, *n*=30 females Mean Age ± SD: 72.9 ± 7.3). Participants from the ADNI 2 databases who presented with MCI were further subdivided in the database into into early-stage MCI (eMCI; *n*=30 males Mean Age ± SD: 73.1 ± 5.2, *n*=30 females Mean Age ± SD: 73.0 ± 8.2), late-stage MCI (lMCI; *n*=29 males Mean Age ± SD: 74.0 ± 8.1, *n*=26 females Mean Age ± SD: 72.0 ± 7.8). A third group of participants from the ADNI 1 database who were not classified into early or late MCI were presumed to be comprised of a broad spectrum of MCI and were therefore treated as a bridge between the eMCI and lMCI groups in the current analyses (MCI; *n*=30 males Mean Age ± SD: 73.1 ± 7.5, *n*=30 females Mean Age ± SD: 73.7 ± 7.2). Neuroimaging data was downloaded from the ADNI data repository August 25, 2023. Ethical approval was obtained by the ADNI investigators at each participating ADNI site, all participants provided written informed consent.

### Transcriptomics

Detailed information describing sample preparation for Xenium spatial transcriptomics and GeoMX Digital Spatial Profiling, as well as the analysis of these transcriptomic techniques, can be found in the Supplemental Methods.

### Electrophysiology

400 μm coronal slices comprising the RSC were obtained in oxygenated ice-cold sucrose/ACSF using the Leica VT1000 S vibratome. Slices were recovered at 30C in ACSF (128 mM NaCl, 10 mM D-glucose, 26 mM NaHCO_3_, 2 mM CaCl_2_, 2 mM MgSO_4_, 3 mM KCl, 1.25 mM NaH_2_PO_4_, pH7.4) saturated with 95% O_2_ and 5% CO_2_. Recordings were performed after a minimum 2-hour recovery. Patch pipettes (2-4 MOhm) contained the following internal patch solution: 120 mM potassium gluconate, 10 mM HEPES, 5 mM KCl, 2 mM MgCl_2_, 4 mM K_2_-ATP, 0.4 mM Na_2_-GTP, 10 mM Na_2_-phosphocreatine, at pH 7.3. Neurons were visualized using the IR-DIC on an Olympus BX51WI microscope. Whole-cell patch clamp recordings were performed in current-clamp and voltage-clamp mode using a MultiClamp 700B amplifier (Molecular Devices). In current-clamp mode, action potentials were elicited by applying 500 ms depolarizing current pulses with a step size of 50 pA. For sIPSC recordings, patch pipettes contained the high Cl^−^ patch solution: 50 mM K-gluconate, 10 mM HEPES, 75 mM KCl, 2 mM MgCl_2_, 4 mM K_2_-ATP, 0.4 mM Na_2_-GTP, 10 mM Na_2_-phosphocreatine, at pH 7.3. sIPSC recordings were performed in voltage clamp mode in the presence of 20 μM AMPA/Kainate receptor antagonist CNQX (Tocris) and 50 μM NMDA receptor antagonist AP5 (Tocris). Data were filtered at 4 kHz and digitized at 20 kHz using Digidata 1550B and Clampex software (Molecular Devices). Recordings were analyzed using pClamp 10.7 or Easy Electrophysiology.

### Behavioral testing

Mice were habituated to handling for 5 days prior to all behavioral tasks. In all tasks, mouse behavior was tracked using ANYmaze Behavioral Tracking Software (Stoelting Co., Wood Dale, IL, USA). Before and after each trial, all apparatuses were cleaned using 70% EtOH.

#### Forced Alternation

Working memory ability was assessed using a Y-maze forced alternation task. The arms of the Y-maze apparatus were 41.5 cm long and 7.5 cm wide. Mice were placed at the end of one arm of the Y maze allowed 5 min to explore the apparatus. During this trial, one of the arms of the Y-maze was blocked. After a 30 min inter-trial interval, mice were returned to the Y-maze and were given 5 min to explore the apparatus without any of the arms being blocked. Time spent in the novel arm of this maze was assessed as a percentage of the total exploratory time, defined as the total time remaining in the task after the animal first leaves the start arm.

#### Contextual Conditioning Task

Contextual conditioning took place in Ugo Basile (Gemonio, Italy) contextual fear conditioning chambers (17 cm X 17 cm) placed inside of sound attenuating cabinets. During the 5 min conditioning trial, mice were allowed to explore the chamber for 2 min prior to the delivery of three shocks (1 mA, 2 s), each separated by an interval of 1 min. Mice were removed from the conditioning chamber 1 min after the final shock and returned to the home cage. Context reintroduction took place 24 h later, during which mice were returned to the conditioned context for a 5 min trial without any shocks. Freezing was used as the primary measure of memory retention and was defined as a complete lack of motion, except for respiration, for at least one second. In cases of baseline differences in freezing during the initial conditioning trials, activity suppression was used to assess memory retention. Activity suppression was calculated as the distance travelled during the first 2 min of the retention trial divided by the total distance travelled during the first two minutes of the retention and conditioning trials(*32*).

### Histology

#### Mouse tissue: perfusion and tissue processing

Ninety minutes following the completion of contextual conditioning tasks, mice were deeply anesthetized with isoflurane and transcardially perfused with 0.1 M phosphate-buffered saline (PBS) and 4% formaldehyde. Brains were extracted and postfixed in 4% formaldehyde at 4 °C for 24 h before cryoprotection in 30% w/v sucrose solution in PBS for two to three days until no longer buoyant. Brains were sectioned 50 μm thick on a cryostat (Leica CM 1950, Concord, ON, Canada) in 12 series. Sections were kept in a buffered antifreeze solution containing 30% ethylene glycol and 20% glycerol in 0.1 M PBS and stored at −20 °C.

#### Immunofluorescent staining

For all immunofluorescent staining, tissue sections were washed three times (10 min per was) in 0.1 M PBS at room temperature. Sections were then incubated at room temperature in primary antibody solution, containing primary antibody, 3% normal goat serum, and 0.3% Triton X-100 (see Table S1 for antibody information, dilutions, and incubation times). Following primary incubation, tissue was washed in 0.1 M PBS (3 x 10 mins) prior to incubation in secondary antibody solution (see Table S1) in 0.1 M PBS at room temperature. Finally, tissues were counterstained with DAPI (20 mins, 1:1000 in 0.1 M PBS) before being washed in 0.1 M PBS (2 x 10 mins) and mounted to glass slides. Slides were coverslipped with PVA-DABCO mounting medium.

#### Human tissue: tissue processing

Human tissue samples previously fixed and stored in formalin fixative, were cryoprotected in 30% w/v sucrose solution in 0.1 M PBS for 3 – 5 days. Tissue was sectioned 50 μm thick on a cryostat (Leica CM 1950, Concord, ON, Canada) in 12 series. Sections were stored at −20 °C in a buffered antifreeze solution containing 30% ethylene glycol and 20% glycerol in 0.1 M PBS.

#### Peroxidase staining

Human tissue sections were stained as previously described(*33*). Briefly, samples were rinsed in 0.1 M tris-buffered saline (TBS) for 10 min prior to a 10 min incubation in BLOXALL (Vector Laboratories, SP-6000) at room temperature. Tissue was then rinsed with TBS (2 X 5 min) before being incubated in a solution of 2.5% normal goat serum in TBS for 20 minutes. After the serum incubation, tissue was transferred to a primary antibody solution with a 1:500 concentration of anti-parvalbumin antibody (Invitrogen, PA1-933), 2.5% normal goat serum, and 0.3% Triton X-100 in 0.1 M TBS at 4 °C. Tissue was then washed with 0.1 M TBS (5 X 5 min) and incubated in ImmPRESS HRP goat anti-rabbit IgG polymer detection kit reagent (Vector Laboratories, MP-74551) for 18 h at 4 °C. Following this incubation, tissue was washed with 0.1 M TBS (5 X 5) and the stain was allowed to develop for 2 minutes using an ImmPACT novaRED Substrate Kit (Vector Laboratories, SK-4805). Tissue sections were then washed for 5 min in tap water, followed by rinses in 0.1M TBS (2 x 5 min). Sections were mounted on glass slides and dried on a slide warmer for 15 minutes and were then dehydrated in two 3-minute incubations in isopropyl alcohol. Slides were coverslipped with VectaMount Express Mounting Medium (H-5700).

### Imaging and image analysis

#### Optical density analyses

The optical density of cytochrome C oxidase and GLUT3 expression was assessed using images collected at 10X magnification (N.A. 0.4) using an OLYMPUS VS120-L100-W slide scanning microscope (Richmond Hill, ON, CA). Brightfield microscopy was used in collecting these images for the cytochrome C oxidase-stained tissue, while TRITC and DAPI filter cubes were used to collect images of the GLUT3-stained tissue. All optical density analyses were conducted using ImageJ.

#### Cellular density analyses

The density of parvalbumin, NeuN, and c-Fos expression density in mouse brain tissue was assessed using images collected at 10X magnification (N.A. 0.4) using an OLYMPUS VS120-L100-W slide scanning microscope (Richmond Hill, ON, CA). In fluorescently stained mouse tissue, labels were segmented from background based on label size and fluorescent intensity using the user-guided machine learning image processing software *Ilastik*. Binary segmented labels were exported from *Ilastik* and their expression density was assessed within the target region based on a tracing of this region in the accompanying DAPI channel image in ImageJ.

In human tissue PV-INs were segmented using the machine-learning based TrueAI software from OLYMPUS. With this segmentation, the machine learning protocol was trained to a subset of manually classified PV-INs and allowed one million iterations to establish a classification algorithm. The accuracy of this algorithm was assessed at 5,000, 10,000, 25,000, 100,000, 250,000, and 500,000 iterations. In mouse and human analyses, all cell counts were normalized based on the area of the target region.

#### Synaptotagmin-2 analyses

Analyses of Syt2 expression were conducted using an OLYMPUS FV3000 confocal microscope equipped with a 60X oil immersion objective (N.A. 1.42). In assessing the broad expression density of Syt2 in the RSC, volumetric scans (60X total magnification; 10 z steps; 0.35 μm z spacing) were collected in this region. Syt2 puncta were segmented from maximum intensity projections of these image stacks using *Ilastik* and their density was assessed using ImageJ. To assess the density of Syt2 puncta on PV-INs and CaMKII+ cells, a 5X zoom was applied for the collection of 20 volumetric scans (300x total magnification; full cells in z dimension; 0.35 μm z spacing) per mouse. Maximum intensity projections were generated from the top half of each volumetric scan and Syt2 puncta were segmented using *Ilastik*. PV+ and CaMKII+ cells were traced using ImageJ and the density of Syt2 puncta was assessed within the bounds of these tracings.

#### Functional connectivity analyses

Brain-wide c-Fos expression was imaged using an OLMYPUS VA120-L100-W slide scanning microscope (Richmond Hill, ON, CA) equipped with a 10X objective (N.A. 0.4). Using *Ilastik*, c-Fos+ cells were segmented to generate binary masks of brain-wide c-Fos expression(*34*). These binary masks were then registered to the Allen Mouse Brain Reference Atlas using a modified protocol building upon *Whole Brain*(*35, 36*). Using custom MATLAB analyses, outputs were organized to yield c-Fos expression densities across 90 neuroanatomical regions (for a detailed list of regions, see Table 2). Regional c-Fos expression density was correlated across groups for all possible combinations of regions, generating correlation matrices of regional co-activation for each group(*37*). From each co-activation matrix, the functional connectivity of the RSC was assessed using calculation derived from the Brain Connectivity Toolbox(*24*). For analyses of binary network characteristics, such as local efficiency and clustering coefficient, co-activation matrices were binarized to adjacency matrices using thresholding criteria of *P* < 0.05 and Pearson’s *R* > 0.8.

#### Validation of viral targeting

Prior to any statistical analyses in the fiber photometry and chemogenetic experiments, the accuracy of virus/optic fiber targeting. For this validation, a series of DAPI-stained tissue sections from each mouse were scanned at 10X magnification (N.A. 0.4) using an OLYMPUS BX-63 epifluorescent microscope (Richmond Hill, ON, CA). The location and spread of the viral injections were then assessed, blind to treatment, relative to the target surgical coordinates using the Allen Mouse Brain Reference Atlas as a reference. All data from any mice showing inadequate viral targeting or expression was excluded from analyses.

### *In vivo* calcium imaging experiments

#### General surgical procedures

Surgeries were conducted under isoflurane delivered via a Somnosuite anaesthetic delivery system (Kent Scientific) drawing pure compressed oxygen. Mice were induced at 5% isoflurane before being transferred to a stereotaxic head frame (Kopf) and maintained at 1-2% isoflurane. Analgesia (Metacam, 5 mg/kg) and fluid support (warmed saline, 0.5 mL) were given at the beginning of the surgery. The scalp was shaved and cleaned with alternating chlorhexidine and 70% ethanol scrubs prior to midline incision, exposing the skull.

Detailed information on miniscope and fiber photometry surgical procedures can be found in the Supplemental Methods.

#### Miniscope recordings

Mice were habituated to being connected to an open-source UCLA v4 Miniscope (https://github.com/Aharoni-Lab/Miniscope-v4) for 3 consecutive days(*38*). Recordings were collected using the Miniscope data acquisition board (DAQ) 3.3 and controlled using the Miniscope QT Software (https://github.com/Aharoni-Lab/Miniscope-DAQ-QT-Software). Recordings occurred throughout the entire duration of the behavioral task and behavior was time locked to the Miniscope signal using timestamp information provided by the DAQ. Behavioral performance was assessed by processing timelocked videos with the open-source software ezTrack(*39*).

#### Photometry recordings

Mice were habituated to being connected to the fiber optic path cord (Doric) for 3 consecutive days. Photometry recordings were collected using a FP3002 Neurophotometrics system controlled by Bonsai software. Prior to the start of each behavioral test, a 5-minute baseline photometry recording was taken in a clean and empty home cage. Recordings occurred throughout the entire duration of the behavioral task and behavior was time locked to the photometry signal through a TTL pulse generated by ANYmaze tracking software (Stoelting) which was integrated into the Neurophotometrics photometry data. Recordings were acquired at 40 FPS, with 470 nm and 415 nm channels being active in alternating frames, resulting in an effective framerate of 20 FPS for each channel. Both the 470 nm channel and the 415 nm isosbestic channel were calibrated to 50 uW using a photometer (Thor Labs PM100D).

#### Analyses

Detailed information on miniscope and fiber photometry surgical analyses can be found in the Supplemental Methods.

### Neuroimaging experiment

Detailed information describing image acquisition protocols, image processing procedures, and functional connectivity analyses can be found in the Supplementary Methods.

### Chemogenetic experiments

#### Viruses

All virus was diluted 1:3 in sterile saline prior to injection. For chemogenetic inhibition experiments, mice received injections of either hM4Di virus or a control virus (pAAV-s5e2-GFP; Addgene #135631-AAV1). The hM4Di virus was derived from Addgene plasmid 83896. The hDlx enhancer element was removed and replaced with the s5e2 regulatory element and packaged as AAV1 by the Hotchkiss Brain Institute Molecular Core Facility. For the experiments with chemogenetic activation, all mice received injections of a hM3Dq virus (pAAV-s5e2-Gq-P2A-dTomato; Addgene #135635-AAV1)(*40*).

#### Surgical procedures

Surgical procedures were conducted as described in the fiber photometry experiments. Virus was injected in four 50 nL pulses at a rate of 10 nL/s with 10 s between pulses at the anterior (AP: −0.9; ML: ± 0.3; DV: 1.2) and posterior (AP: −2.2; ML: ± 0.3; DV: 1.0) portions of the RSC. The pipette was left in the target for 5 minutes following the final pulse before being slowly raised out of the brain. The incision was then closed with suture material and mice were removed from the stereotaxic frame. Mice were allowed to recover for 2 weeks while receiving additional Metacam doses for 3 days after surgery.

#### DCZ preparation and administration

Deschloroclozapine dihydrochloride (DCZ; HelloBio HB126-25mg) was diluted in saline for a working dose of 3 μg/kg (0.3 mg/mL, inj. at 0.01 mL/g)(*41*). For chemogenetic inhibition experiments, all mice received DCZ treatment 15 minutes prior to the start of each behavioral test. For chemogenetic activation experiments, mice received either DCZ treatment or saline injection (0.01 mL/g) 15 minutes prior to the start of each behavioral test.

### Statistical analysis and data visualization

All statistical analyses were performed in Prism (GraphPad Software, Version 9.4.0) or with standard MATLAB functions and statistical tests. Independent t-tests; three-, two-, and one-way ANOVAs; and Pearson’s Correlations were performed. Data in graphs is presented as mean ± s.e.m. Hypothesis testing was complemented by estimation statistics for each comparison using estimationstats.com(*42*). For these estimation statistics, the effect size (Cohen’s d) was calculated using a bootstrap sampling distribution with 5000 resamples along with a 95% confidence interval (CI; bias-corrected and accelerated). Data distribution was assumed to be normal, but this was not formally tested. Statistical methods were not used to predetermine study sizes but were based on similar experiments previously published. Experimenters were blinded to the genotype and sex of the animals during all analyses. Experimenters were blinded to all identifying features when analyzing human tissue samples. All statistical comparisons and outputs are included in Tables S3 and S4. All plots were generated in Prism or MATLAB. Circle plots were generated using the circularGraph MATLAB function. Some of the schematics were generated using images from BioRender. All figures were compiled in Adobe Photoshop.

## List of Supplementary Materials

Supplemental Methods

Fig. S1 to S5

Table S1 to S3

## Acknowledgments

Funding for this study was provided by an Alzheimer’s Society Research Program (ASRP) New investigator Grant (#21-05), a Canadian Foundation for Innovation (CFI) Grant (#38160), and a Women’s Brain Health Initiative Grant in partnership with Brain Canada (#5542) to J.R.E. as well as an Alzheimer’s Association Grant in partnership with Brain Canada (AARG-22-917644) to D.S. D.J.T. received a doctoral fellowship from NSERC (PGS D). Y.R. received doctoral awards from the Hotchkiss Brain Institute and Alzheimer’s Society Research Program (ASRP). Transcriptomic analyses were performed with assistance from the Applied Spatial Omics Centre. The znp-1 monoclonal antibody (AB_2315626) developed by Bill Trevarrow and colleagues was obtained from the Developmental Studies Hybridoma Bank, created by the NICHD of the NIH and maintained by the University of Iowa, Department of Biological Sciences, Iowa City, IA, USA. AAV1-s5e2-jGCaMP6f (Addgene viral prep # 135632-AAV9; http://n2t.net/addgene:135632; RRID:Addgene_135632), pAAV-s5e2-GFP (Addgene viral prep # 135631-AAV1; http://n2t.net/addgene:135631; RRID:Addgene_135631), and pAAV-s5e2-Gq-P2A-dTomato (Addgene viral prep # 135635-AAV9; http://n2t.net/addgene:135635; RRID:Addgene_135635) were gifts from Jordane Dimidschstein. AAV-hDlx-GiDREADD-dTomato-Fishell-5 was a gift from Gordon Fishell (Addgene plasmid # 83896; http://n2t.net/addgene:83896; RRID:Addgene_83896). We acknowledge the Hotchkiss Brain Institute Advanced Microscopy Platform and the Cumming School of Medicine for support and use of the Olympus VS120-L100-W slide scanning microscope. We acknowledge Dr. Frank Visser and the Hotchkiss Brain Institute Molecular and Cellular Biology Core Facility for generating the s5e2-Gi-P2A-dTomato viral prep. Neuroimaging data used in the preparation of this manuscript were obtained from the Alzheimer’s Disease Neuroimaging Initiative (ADNI) database, which was funded by the National Institutes of Health (Grant U01 AG024904) and DOD ADNI (Department of Defense award number W81XWH-12-2-0012). ADNI is funded by the National Institute on Aging, the National Institute of Biomedical Imaging and Bioengineering, and through generous contributions from the following: AbbVie, Alzheimer’s Association; Alzheimer’s Drug Discovery Foundation; Araclon Biotech; BioClinica, Inc.; Biogen; Bristol-Myers Squibb Company; CereSpir, Inc.; Cogstate; Eisai Inc.; Elan Pharmaceuticals, Inc.; Eli Lilly and Company; EuroImmun; F. Hoffmann-La Roche Ltd and its affiliated company Genentech, Inc.; Fujirebio; GE Healthcare; IXICO Ltd.; Janssen Alzheimer Immunotherapy Research & Development, LLC.; Johnson & Johnson Pharmaceutical Research & Development LLC.; Lumosity; Lundbeck; Merck & Co., Inc.; Meso Scale Diagnostics, LLC.; NeuroRx Research; Neurotrack Technologies; Novartis Pharmaceuticals Corporation; Pfizer Inc.; Piramal Imaging; Servier; Takeda Pharmaceutical Company; and Transition Therapeutics. The Canadian Institutes of Health Research is providing funds to support ADNI clinical sites in Canada. Private sector contributions are facilitated by the Foundation for the National Institutes of Health (www.fnih.org). The grantee organization is the Northern California Institute for Research and Education, and the study is coordinated by the Alzheimer’s Therapeutic Research Institute at the University of Southern California. ADNI data are disseminated by the Laboratory for Neuro Imaging at the University of Southern California.

## Author contributions

Conceptualization: DJT, DS, JRE

Methodology: DJT, YR, BA, HS, DS, JRE

Investigation: DJT, LAMG, DS, JRE Visualization: DJT, DS, JRE

Supervision: DS, JRE Writing—original draft: DJT, JRE

Writing—review & editing: DJT, YR, BA, HS, LAMG, DS, JRE

## Data and materials availability

Neuroimaging data used in this study are available upon request and approval from the ADNI Data and Publications Committee. Instructions for making a request can be found at https://adni.loni.usc.edu/data-samples/access-data/. The members of the Alzheimer’s Disease Neuroimaging Initiative (ADNI) can be found online at http://adni.loni.usc.edu/wp-content/uploads/how_to_apply/ADNI_Acknowledgement_List.pdf.

Sequencing data will be made available through the NCBI Gene Expression Omnibus data repository upon publication.

Code used for the analysis of the transcriptomic data reported in this manuscript along with instructions on how to use these analyses will be made publicly available upon publication.

All other analysis code used to generate the results reported in the manuscript, instructions on how to use these analyses, and the data that support the findings of this study have been made publicly available at the following GitHub repository: https://github.com/dterstege/PublicationRepo/tree/main/Terstege2023A. These files have also been made available through https://doi.org/10.5281/zenodo.13769574.

Further information and requests for resources and reagents should be directed to and will be fulfilled by the lead contact, Jonathan R. Epp.

