## Supplementary material for "Impaired parvalbumin interneurons in the retrosplenial cortex as the cause of sex-dependent vulnerability in Alzheimer’s disease": Terstege2024_SupplementalFigures.pdf

**SUPPLEMENTAL FIGURES**

Supplemental Figure 1

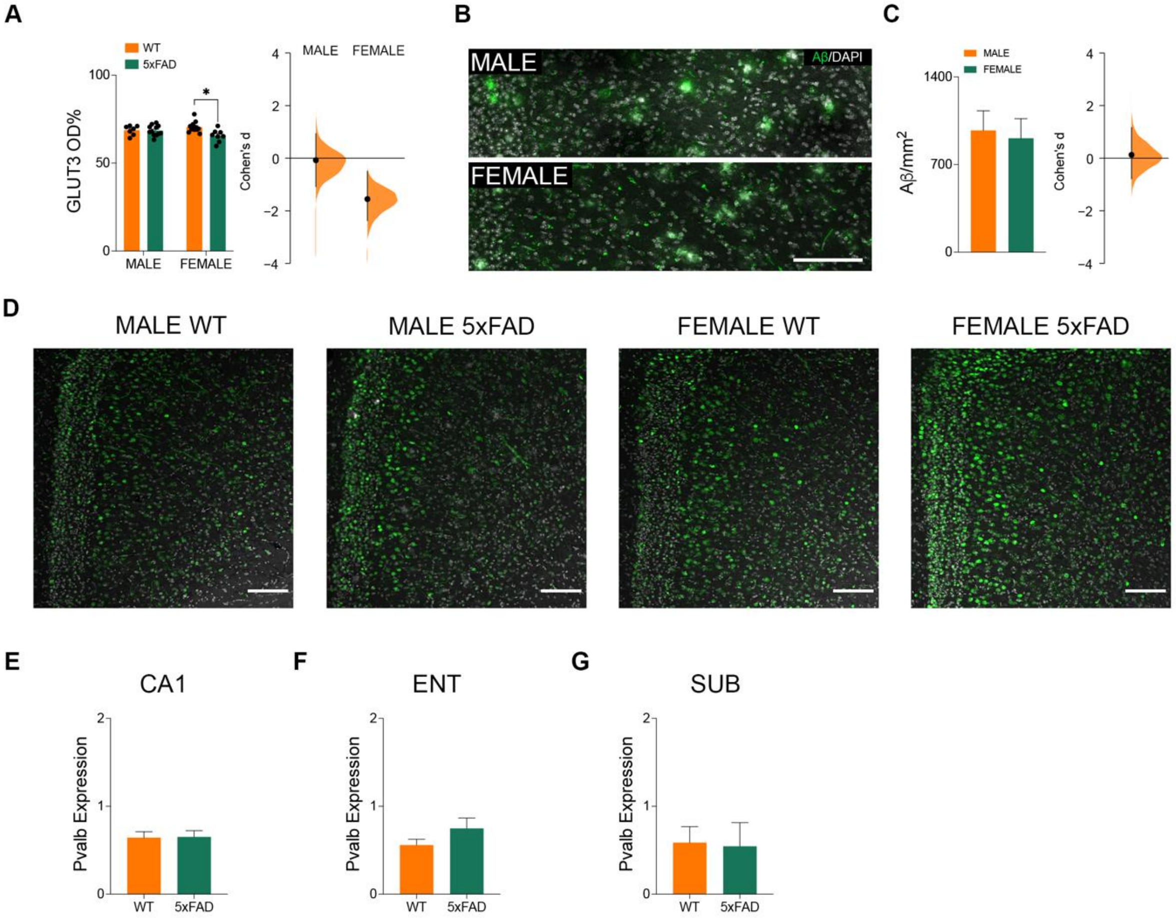


**Fig. S1: No sex differences in RSC amyloid accumulation in 5xFAD mice.** (**A**) Optical density of GLUT3 staining in the RSC. *P*=0.0055, Two-way ANOVA, Tukey's multiple comparison test. Cohen's D effect size between WT and 5xFAD for males (-0.0773) and females (-1.55). (**B**) Amyloid (green) and DAPI (greyscale) staining in the RSC of male and female 5xFAD mice. Scale bar represents 100 µm. (**C**) Density of amyloid aggregation in the RSC of male and female 5xFAD mice. *P*=0.7887, Two-sample t test. Effect size difference in the density of amyloid aggregation in the RSC between male and female 5xFAD mice (Cohen's D=0.126). (**D**) c-Fos (green) and DAPI (greyscale) in the RSC of male and female WT and 5xFAD mice. Scale bars represent 50µm. Normalized *Pvalb* expression in the (**E**) CA1, (**F**) ENT, and (**G**) SUB of 5xFAD mice. Data represent mean ± SEM. All statistical comparisons have been provided in Table S3.

Supplemental Figure 2

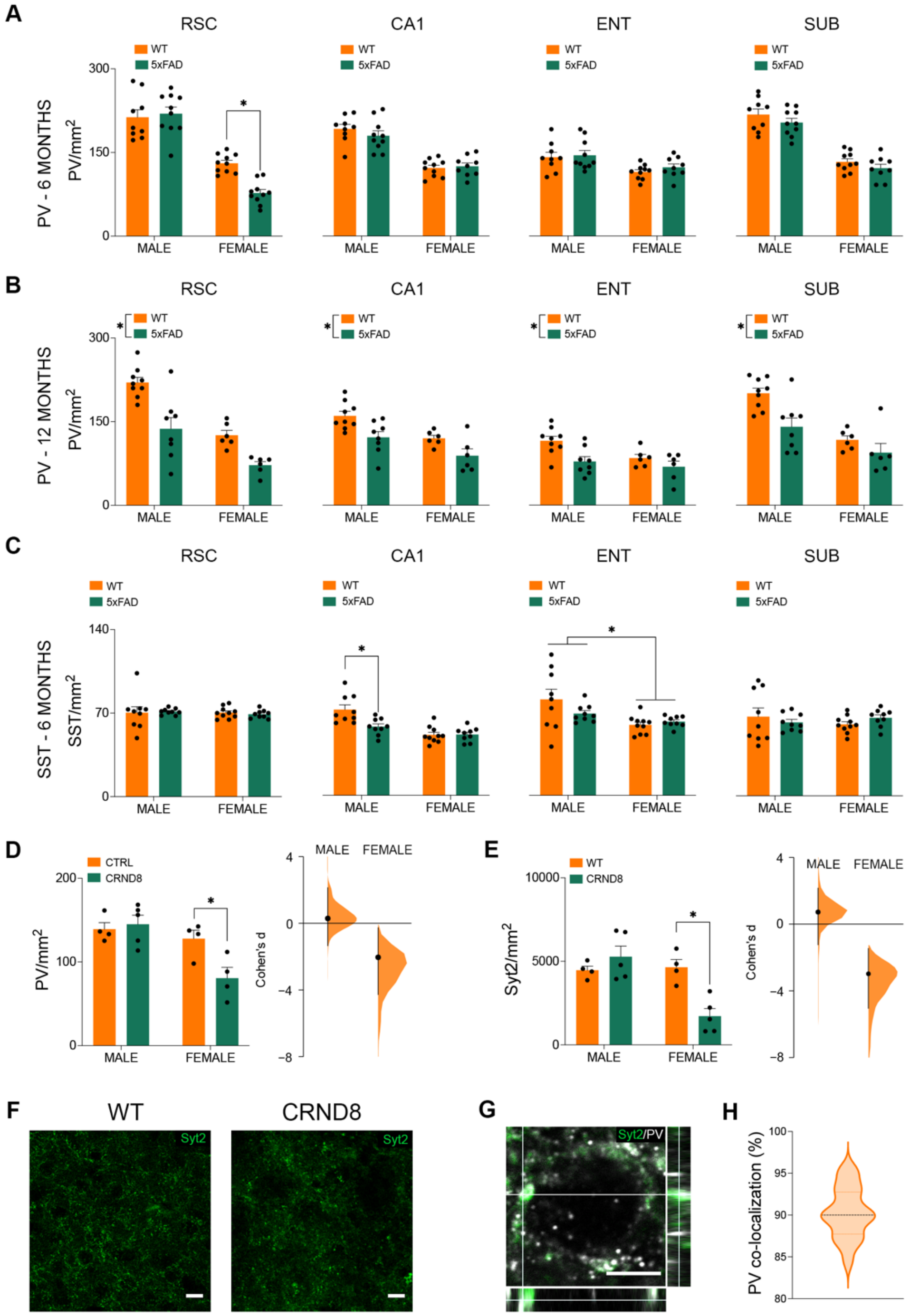

**Fig. S2: Early disruption of PV-INs in the female RSC is cell-type and region-specific.** (**A**) Density of PV-INs in 6-month-old male and female WT and 5xFAD mice. RSC; Female WT VS Female 5XFAD, *P*=0.0019, Two-way ANOVA with Tukey's multiple comparison test. (**B**) Density of PV-INs in 12-month-old male and female WT and 5xFAD mice. RSC; WT vs 5xFAD, *P*<0.0001, Two-way ANOVA. CA1; WT vs 5xFAD, *P*=0.0013, Two-way ANOVA. ENT; WT vs 5xFAD, *P*=0.0058, Two-way ANOVA. SUB; WT vs 5xFAD, *P*=0.0039, Two-way ANOVA. (**C**) Density of SST+ cells in 6-month-old male and female WT and 5xFAD mice. CA1; Male WT VS Male 5XFAD, *P*=0.0045, Two-way ANOVA with Tukey's multiple comparison test. ENT; Male vs Female, *P*=0.0040, Two-way ANOVA. (**D**) Density of PV-INs in the RSC of male and female WT and CRND8 mice. *P*=0.0413, Two-way ANOVA with Tukey's multiple comparison test. Effect size difference in Syt2 density between WT and CRND8 for males (Cohen's D=0.286) and females (Cohen's D=-2.04). (**E**) Density of Syt2 puncta in the RSC of male and female WT and CRND8 mice. *P*=0.0049, Two-way ANOVA with Tukey's multiple comparison test. Effect size difference in Syt2 density between WT and CRND8 for males (Cohen's D=0.725) and females (Cohen's D=-2.98). (**F**) Syt2 staining in the RSC of WT and CRND8 mice. Scale bars represent 10 µm. (**G**) Syt2 (green) and PV (greyscale) staining. Scale bar represents 5 µm. (**H**) Distribution of the percentage of Syt2 puncta co-localized with PV staining across 20 cells. These labels co-localized with a mean of 90.27% ± 3.052% SEM. Data represent mean ± SEM. All statistical comparisons have been provided in Table S3.

Supplemental figure 3

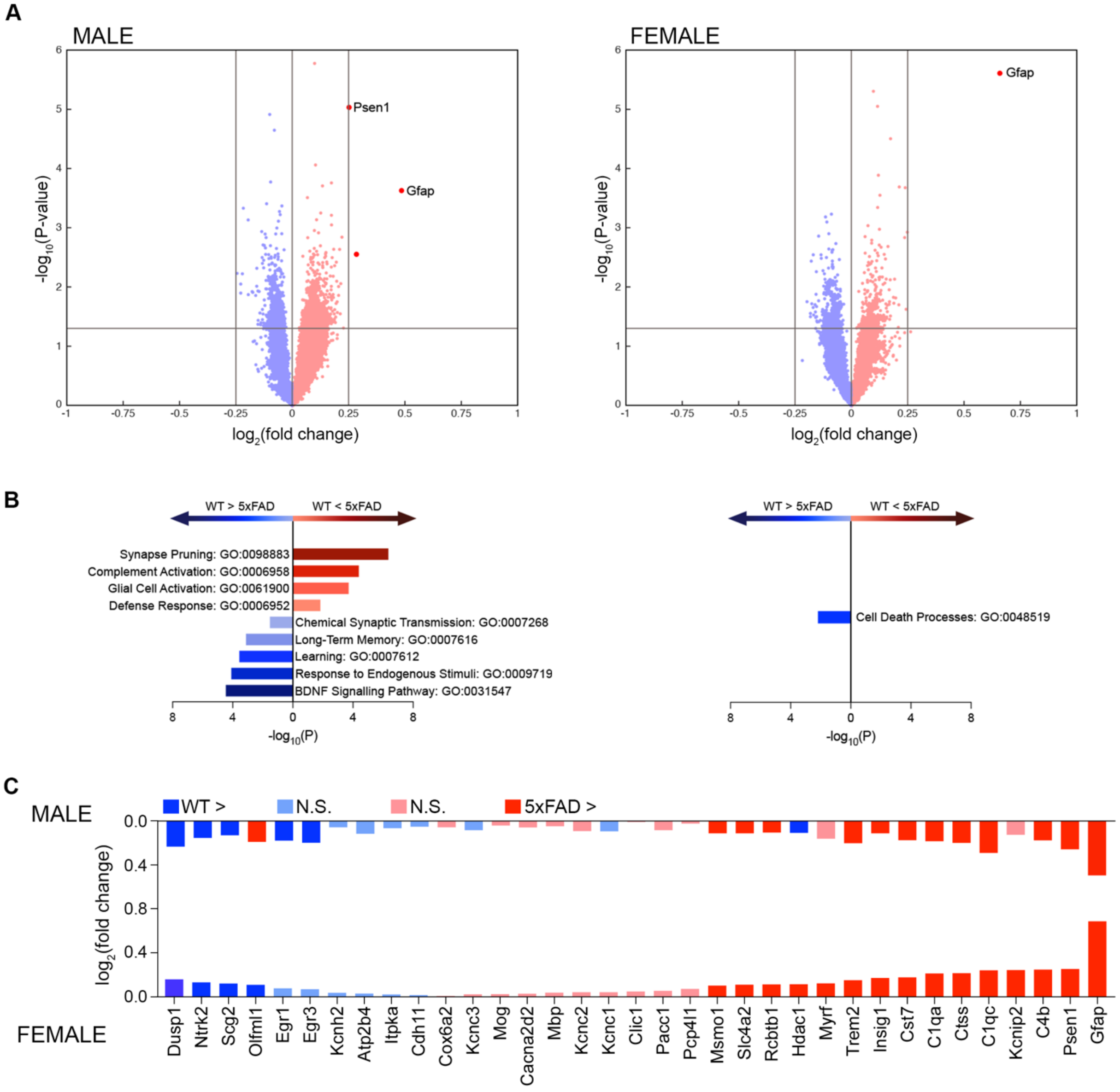


**Fig. S3: Transcriptomic profiles of non-PV-IN, NeuN positive, neurons.** (**A**) Volcano plots of differentially expressed genes (DEG) across the RSC of 5xFAD and WT mice. Dark blue, significantly downregulated in 5xFAD; dark red, significantly upregulated in 5xFAD; light colors, non-significant differential expression. FDR-corrected (95%) Wilcoxon rank-sum test. (**B**) Top GO terms (by fold enrichment) identified by examining the top and bottom 30 DEG in PV-INs between male (left) and female (right) 5xFAD and WT mice. Blue, significantly downregulated gene clusters in 5xFAD relative to WT; red, significantly upregulated gene clusters in 5xFAD relative to WT. (**C**) $Log_2$ fold changes observed between 5xFAD and WT mice in several selected genes in males (top) and females (bottom). Dark blue, significantly downregulated in 5xFAD; dark red, significantly upregulated in 5xFAD; light colors, non-significant differential expression. All statistical comparisons have been provided in Table S3.

### Supplemental Figure 4

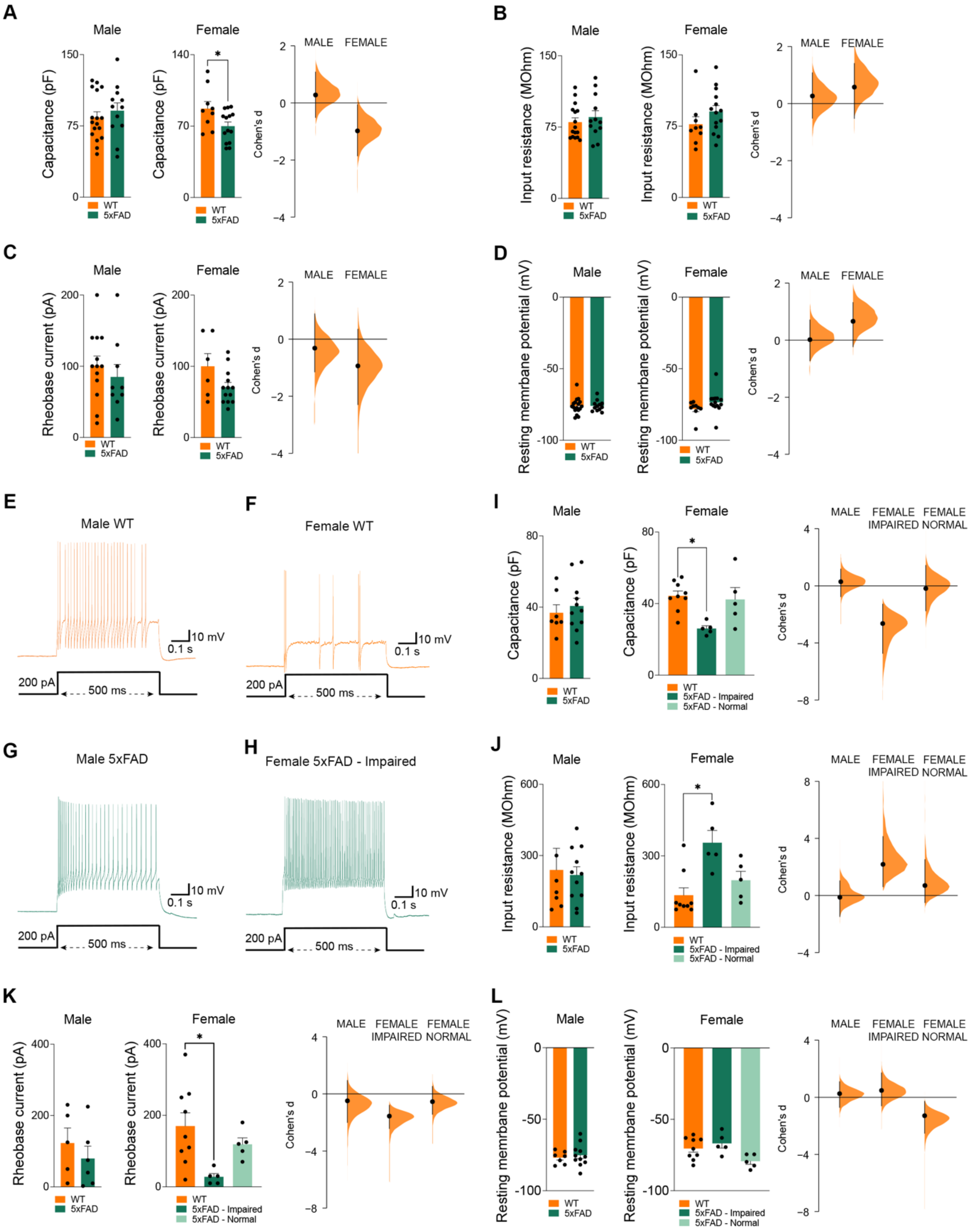

**Fig. S4: Impaired passive membrane properties of PV-INs but not pyramidal cells in female 5xFAD mice.** (**A**) Pyramidal cell membrane capacitance (pF) in WT and 5xFAD mice. *P*=0.0320, Two-sample t test. Effect size difference in membrane capacitance between WT and 5xFAD for males (Cohen's D=0.274) and females (Cohen's D=-0.982). (**B**) Pyramidal cell input resistance (MOhm) in WT and 5xFAD mice. Effect size difference in input resistance between WT and 5xFAD for males (Cohen's D=0.266) and females (Cohen's D=0.577). (**C**) Pyramidal cell rheobase current (pA) in WT and 5xFAD mice. Effect size difference in rheobase current between WT and 5xFAD for males (Cohen's D=-0.314) and females (Cohen's D=-0.938). (**D**) Pyramidal cell resting membrane potential (mV) in WT and 5xFAD mice. Effect size difference in resting membrane potential between WT and 5xFAD for males (Cohen's D=0.0142) and females (Cohen's D=0.659). Current-clamp traces of PV-INs in (**E**) male WT, (**F**) male 5xFAD, (**G**) female WT, and (**H**) female 5xFAD – Impaired groups in response to a 200 pA depolarization step. (**I**) PV-IN membrane capacitance (pF) in WT and 5xFAD mice. *P*=0.0065, One-way ANOVA, Dunnett's multiple comparison test. Effect size difference in membrane capacitance between WT and 5xFAD for males (Cohen's D=0.286) and females (Impaired: Cohen's D=-2.64 Normal: Cohen's D=-0.187). (**J**) PV-IN input resistance (MOhm) in WT and 5xFAD mice. *P*=0.0017, One-way ANOVA, Dunnett's multiple comparison test. Effect size difference in input resistance between WT and 5xFAD for males (Cohen's D=-0.124) and females (Impaired: Cohen's D=2.18 Normal: Cohen's D=0.700). (**K**) PV-IN rheobase current (pA) in WT and 5xFAD mice. *P*=0.0125, One-way ANOVA, Dunnett's multiple comparison test. Effect size difference in rheobase current between WT and 5xFAD for males (Cohen's D=-0.489) and females (Impaired: Cohen's D=-1.56 Normal: Cohen's D=-0.546). (**L**) PV-IN resting membrane potential (mV) in WT and 5xFAD mice. Effect size difference in resting membrane potential between WT and 5xFAD for males (Cohen's D=0.256) and females (Impaired: Cohen's D=0.474 Normal: Cohen's D=-1.29). Data represent mean ± SEM. All statistical comparisons have been provided in Table S3.

Supplemental Figure 5

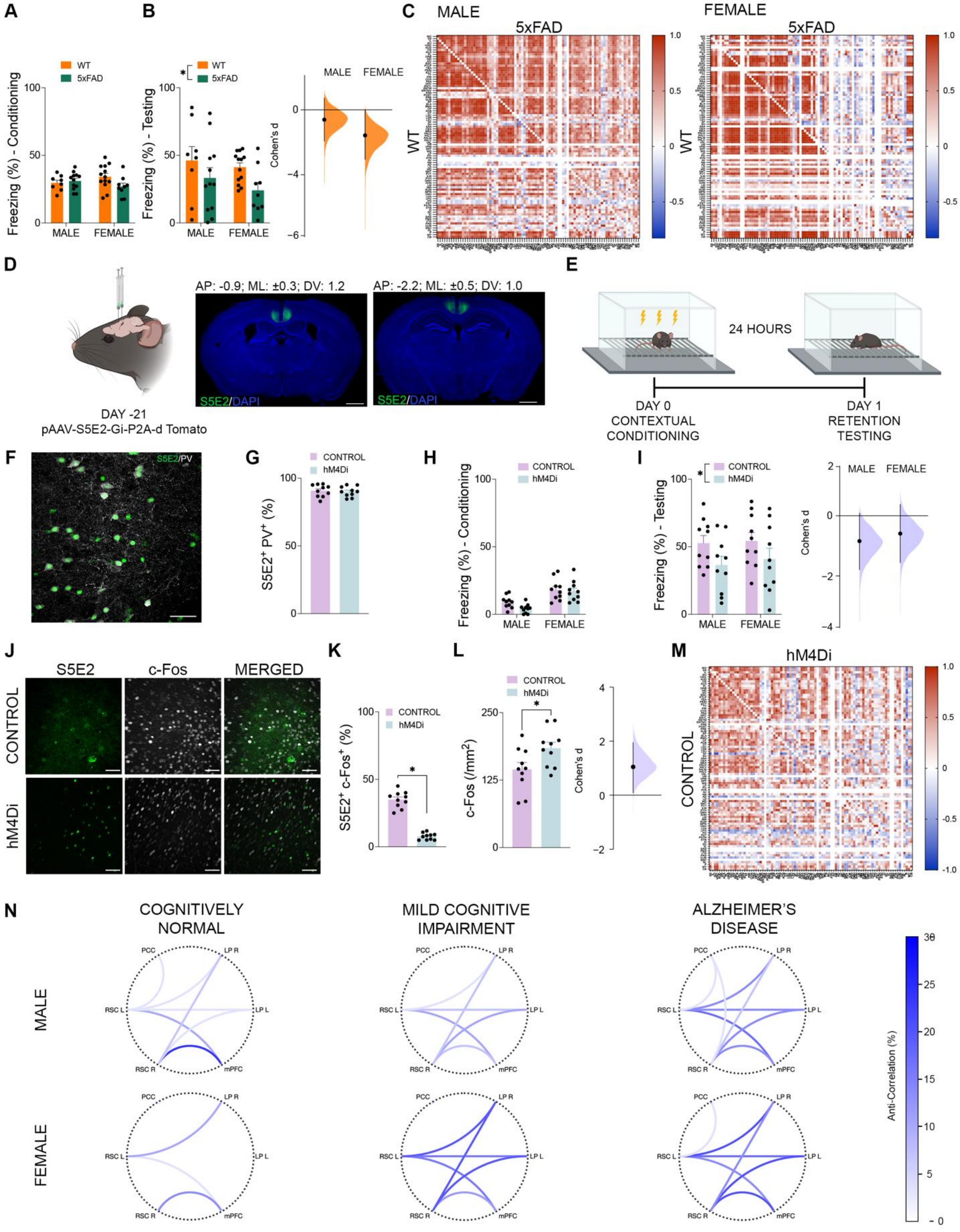

**Fig. S5: Pattern of altered RSC functional connectivity in female 5xFAD mice, healthy mice with dampened RSC PV-IN activity, and in clinical AD progression.** (**A**) Percentage of time spent freezing during conditioning by WT and 5xFAD. (**B**) Percentage of time spent freezing upon reintroduction to the conditioned context by WT and 5xFAD mice. $P$=0.0324, Two-way ANOVA. Effect size difference in percentage of time spent freezing between WT and 5xFAD for males (Cohen's D=-0.469) and females (Cohen's D=-1.22). (**C**) Correlation matrices showing the co-activation of 90 brain regions based on c-Fos expression density in male mice. The WT group is shown on the bottom left diagonal with the 5xFAD group shown on the top right diagonal. (**D**) Schematic showing pAAV-S5E2-Gi-P2A-d Tomato (hM4Di) or pAAV-S5E2-GFP (control) virus injection site and representative photomicrographs of S5E2 (green) viral labelling and DAPI staining (blue). Scale bars represent 1 mm. (**E**) Schematic outlining the contextual fear conditioning task. (**F**) S5E2 (green) labelling and PV (greyscale) staining. Scale bar represents 50 μm. (**G**) Percentage of S5E2+ cells which where colocalized with PV in control and hM4Di treated mice. Together, these labels co-localized with a mean of 90.42% ± 4.058%. (**H**) Percentage of time spent freezing during contextual conditioning by mice injected with hM4Di or control virus. (**I**) Percentage of time spent freezing upon reintroduction to the conditioned context by mice injected with hM4Di or control virus. $P$=0.0317, Two-way ANOVA. Effect size difference in percentage of time spent freezing between control and hM4Di for males (Cohen's D=-0.844) and females (Cohen's D=-0.594). (**J**) S5E2 (green) viral labelling and c-Fos (greyscale) staining with control virus and hM4Di treatment. Scale bars represent 50 μm. (**K**) Percentage of S5E2+ cells which were colocalized with c-Fos with control virus and hM4Di treatment. $P$<0.0001, Two-sample t test. (**L**) Density of c-Fos+ cells in the RSC in mice injected with control virus or S5E2-hM4Di. $P$=0.0308. Effect size difference in RSC c-Fos density between control and hM4Di (Cohen's D=1.05). (**M**) Correlation matrices showing the co-activation of 90 brain regions based on c-Fos expression density. The control virus group is show on the bottom left diagonal with the hM4Di group shown on the top right diagonal. (**N**) Percentage of subjects with anti-correlated resting-state functional connectivity between specific regions within the default mode network. Data represent mean ± SEM. All statistical comparisons have been provided in Table S3.
