## Supplementary material for "Impaired parvalbumin interneurons in the retrosplenial cortex as the cause of sex-dependent vulnerability in Alzheimer’s disease": Terstege2024_SupplementalMethods.pdf

### SUPPLEMENTAL METHODS

#### Transcriptomics:

*Xenium spatial transcriptomics*:

Fixed, paraffin-embedded brains from 6-month-old WT and 5xFAD mice were hemisectioned at thickness of 5 µm and prepared according to recommendations in the Xenium *In Situ* for FFPE (Formalin Fixed & Paraffin Embedded) – Tissue Preparation Guide (10X Genomics, CG000578). Briefly, sections from female WT (*n*=3) and 5xFAD (*n*=3) were mounted within the sample frame of the Xenium slide, deparaffinized, and permeabilize to make the mRNA accessible. Following the Xenium *In Situ* Gene Expression protocol (10X Genomics, CG000582), hybridization was performed overnight at 50 °C using the pre-designed 247-probe Xenium Mouse Brain Gene Expression panel (10X Genomics, Cat#1000462), which includes 2 probes which serve as negative controls. Each of these circularizable DNA probes contains two regions that hybridize to the target RNA and a third region that encodes a gene-specific barcode. Unhybridized probes were removed by washing the slide. Remaining hybridized probes were annealed to their specific mRNA targets through a 2 h ligation step at 37 °C. To enzymatically amplify the signal, samples underwent rolling circle amplification PCR for 2 h at 37 °C. Slides were then washed again and background fluorescence was chemically quenched and a DAPI stain was added to provide contrast for subregion identification.

All instrument-specific reagents and buffers were prepared according to the Xenium Analyzer User Guide (10X Genomics, CG000584). The Xenium slide was loaded onto the Xenium Analyzer platform along with the appropriate reagents and buffers for high throughput, automated *in situ* analysis. A simple preliminary scan of the tissue sections allowed for the identification of regions of interest covering the entire sections, which were selected in the Xenium Analyzer software. With regions identified, slides underwent iterative rounds of image acquisition wherein fluorescently labelled probe-specific oligonucleotides are hybridized to the amplified probe sequence, imaged, and then gently removed. This allowed for the generation of optical signatures for each specific barcode, which would later be converted into gene identities. The Xenium Mouse Brain predesigned panel (version 1.1) comprised 247 genes, and we used the predesigned panel without an add-on panel. Only transcripts with a Xenium Quality Value >20 were retained. Images were acquired with an XY localization precision of <50nm and a Z step size of 0.75 µm. Stitching these images back together created a spatial map of the transcripts across entire tissue sections. Data were processed on-premises using the standard 10X Genomics Xenium workflow to produce cell-by-gene and transcript-by-location matrices.

*Xenium analyses*:

Xenium data was loaded into R (v4.3.1) for analyses using Seurat package (v5.1.0). During preprocessing and quality control procedures, any transcripts which were not assigned to cells identified through the DAPI channel were excluded. Under-segmented delineations, wherein the transcripts from multiple DAPI-positive nuclei were collected as one, were identified using Grubb's test and excluded. Cells with few (<10) detected transcripts and high background signals (proportion of negative probes relative to the total number of probes expressed in the cell > 0.05) were also excluded from these analyses. As per 10X Genomics recommendations, a phred-like quality score threshold of 20 was used to exclude low quality readings.

Following preprocessing and quality control measures, subsets of the transcriptomic expression profiles were generated based on ROIs drawn at the RSC, SUB, ENT, and CA1. Within these ROIs, differential gene expression analyses were conducted using the 10X Genomics enumerated differential expression analysis tools. Using Wilcoxon signed-rank test, the log fold change in gene expression was compared between WT and 5xFAD groups at each of these subregions, with $p$ values adjusted for multiple comparisons.

*GeoMX digital spatial profiling*:

Fixed, paraffin-embedded brains from 6-month-old WT and 5xFAD mice were hemisectioned at thickness of 5 μm and prepared for GeoMX Digital Spatial Profiling (DSP)*(43)*. Briefly, unstained sections containing the RSC of male WT ($n$=4), male 5xFAD ($n$=4), female WT ($n$=3), and female 5xFAD ($n$=4) mice were mounted to slides for the GeoMX DSP (NanoString, Inc., v2.5) assay. Slides were baked for 3 h at 60 °C, deparaffinized with xylene, and then rehydrated through graded ethanol washes. Transcript targets were exposed by incubating the slides in pH 9 Tris-EDTA buffer followed by a solution of 0.1 μg/mL proteinase K. *In situ* hybridizations were performed with the GeoMX Mouse Whole Transcriptome Atlas (20,175 total targets, including 210 negative probes targeting sequences which are not present in the genome) according to the manufacturer's instructions (NanoString, Inc., MAN-10150). Probes were added to each slide in a hybridization chamber, covered with a coverslip, and incubated overnight at 37 °C. 24 h later, slides were washed with a solution of SSC/50% formamide and blocked in Buffer W from the GeoMX DSP assay kit. Slides were then incubated for 1 h in a solution of 1:100 Alexa Fluor 488 conjugated anti-NeuN (Sigma-Aldrich; ABN78A4), 1:100 Alexa Fluor 594 (Abcam; ab269822) conjugated in-house to an anti-PV antibody (Invitrogen; PA1-933), and 1:50 SYTO 83 (Thermo Fisher, S11364) in Buffer W. Slides were briefly washed in SSC to remove any excess markers and then immediately scanned at 20X using the GeoMX Digital Spatial Profiler platform.

Areas of illumination (AOI; 1-2 AOI per section) were drawn onto each section to cover the RSC. Within each AOI, PV+ and NeuN+ labels were segmented from background. Following these initialization procedures, the GeoMX Digital Spatial Profiler platform photocleaves the UV-cleavable barcode linker of the bound RNA probes from the nuclei of all PV+ cell in the AOI and deposits the released oligonucleotides into a well in the DSP collection plate. Oligonucleotides from the nuclei of all NeuN+ cells in the AOI are then retrieved and deposited in a separate well of the DSP collection plate using the same procedures. This process was repeated at each AOI, with the nuclei from each cell type of each AOI being deposited in separate wells of the DSP collection plate.

For library preparation, Illumina i5 and i7 dual-indexing primers were added to the oligonucleotide tags during polymerase chain reaction (PCR) to uniquely index each AOI. AMPure XP beads (Beckman Coulter) were used for PCR purification. Library concentration was measured using a Qubit fluorometer (Thermo Fisher Scientific), and quality was assessed using a Bioanalyzer (Agilent Technologies). For sequencing of whole transcriptome analysis libraries, the target sequencing depth was 100 count/$\mu m^2$. Sequencing was performed on an Illumina NextSeq 2000 (Illumina Inc.) and FASTQ files were processed into gene count data for each AOI using the NanoString GeoMX NGS Pipeline (NanoString, Inc., MAN-10153).

*GeoMX analyses*:

Initial quality control of the GeoMX data was implemented on a label-by-label basis by assessing the quality of the isolated segments based on recommended thresholds for the percentage of trimmed reads (>80%), the percentage of stitched reads (>80%), the percentage of aligned reads (>75%), the sequencing saturation (>50%), the area of the AOI (>1000μm$^2$), and a minimum number of nuclei (>20) (NanoString, Inc. MAN-10153-01). After assessing the quality of the segments, low performance probes were identified by dividing the geometric mean of a single probe count across all samples against the geometric mean of all the probe counts for that gene. Samples displaying non-specific binding were then identified by defining the limit of quantification as 2 standard deviations above the geometric mean of the negative probes for each sample. AOIs would have been removed from analyses if fewer than 5% of genes were detected above the limit of quantification. Following quality control, 8,113 genetic targets remained.

The raw count matrix was normalized using variance stabilizing transformation (DESeq2)*(44)*. Differential expression within PV+ and NeuN+ cells was calculated across groups for each gene using a linear mixed effect model with the GeoMX Tools R package (v3.4.0). Genes targets with a $p$ value <0.05 and a log fold change greater than 1.0 were considered to be differentially expressed. Volcano plots were generated using the MATLAB 'volcanoplot' function from the RobkaiXplorer toolbox*(45)*. Gene ontology (GO) analyses were performed using g:Profiler*(46)*. FDR-based $p$ value correction was used for all analyses.

#### *In vivo* calcium imaging experiments:

*Miniscope surgical procedures*:

A robotic stereotaxic manipulator paired with stereodrive software (Neurostar) was used to drill a 1 mm craniotomy. Superficial cortical tissue was carefully aspirated (AP: -2.7; ML: -0.6; DV: 0.6) prior to virus injection via a glass infusion needle attached to a Nanoject III infusion system (Drummond Scientific). Virus (AAV1-s5e2-jGCaMP6f; Addgene #135632-AAV1; diluted 1:3 in saline) was injected into the RSC (AP: -2.7; ML: -0.6; DV: 0.8 and 1.2) in six sets of 50 nL pulses at each target depth at a rate of 10 nL/s with 10 s between pulses. The pipette was left at each target depth for 5 minutes following the final pulse to allow for diffusion of the virus before being raised slowly. Following virus injection, a 1 mm diameter GRIN lens (Go!Foton; 3.758 mm length, 0.433 pitch at 550 nm, 0.2 mm working distance) was lowered into the RSC (AP: -2.7; ML: -0.6; DV: 0.8). The GRIN lens was secured to the skull by applying a thin layer of superglue to the exposed surface of the skull around the base on the implant. This layer was then covered with black, opaque, dental acrylic. To protect the lens, the tip of a small PCR tube was glued to the dental cement, once dry. Four weeks later, the protective tube was removed, and excess dental cement was cleared. Using a Miniscope to visually guide the placement, a V4 Miniscope baseplate was attached to the skull. With the baseplate in position, it was secured to with dental cement to form a new headcap. Once this headcap was set, mice were removed from the stereotaxic frame. Mice were allowed to recover for 2 weeks while receiving additional Metacam doses for 3 days after surgery.

*Miniscope analysis*:

Calcium transients were analyzed using the open-source calcium imaging data processing pipeline, Min1pipe*(47)*. Briefly, videos were preprocessed to enhance the neural signals through anisotropic diffusion and morphology opening. Movement correction was then applied to further clean the signal. Cells were delineated using automated seed detection algorithms built into Min1pipe and then refined to remove any imperfect segmentations. From each cell, normalized

ΔF/F traces were extracted and aligned to the timestamps of the zone positional data using nearest neighbor approximation. For analyses of peaks, these were defined as 1 standard deviation above median value of the ΔF/F trace.

*Fiber photometry surgical procedures*:

A robotic stereotaxic manipulator paired with stereodrive software (Neurostar) was used to drill burr holes prior to virus injection via a glass infusion needle attached to a Nanoject III infusion system (Drummond Scientific). Virus (AAV1-s5e2-jGCaMP6f; Addgene #135632-AAV1; diluted 1:3 in saline) was injected into the RSC (AP: -3.1; ML: -0.5; DV: 1.1) in six sets of 50 nL pulses at a rate of 10 nL/s with 10 s between pulses. The pipette was left at the target depth for 5 minutes following the final pulse to allow for diffusion of the virus before being raised slowly. Following virus injection, a fiber optic implant (N.A. 0.37, core 200 μm, 2.0 mm length) was lowered into the RSC at the injection site (AP: -3.1; ML: -0.5; DV: 1.0). Implants were secured to the skull by applying a thin layer of superglue to the exposed surface of the skull around the base on the implant. This layer was then covered with black, opaque, dental acrylic to form a headcap. Once the headcap was set, the incision was closed with suture material and mice were removed from the stereotaxic frame. Mice were allowed to recover for 2 weeks while receiving additional Metacam doses for 3 days after surgery.

*Photometry analysis*:

Photometry recordings were analyzed using custom MATLAB scripts*(48)*. Behavioral information recorded using ANYmaze was integrated into the analysis using a nearest neighbour approach, by indexing through the photometry data and aligning behavioral data based on the minimum difference between pairs of timestamps. Data from the isosbestic 415 nm channel was fit to a biexponential decay to correct for photobleaching. The resulting vector was used to linearly scale calcium-dependent data collected using the 470 nm channel. To calculate ΔF, these linearly scaled calcium-dependent data were subtracted from the raw unprocessed calcium-dependent data. The resulting values were then divided by the linearly scaled calcium-dependent data to generate a ΔF/F trace.

**Neuroimaging experiment:**

*Image acquisition:*

Participants were scanned at multiple sites equipped with 3.0 Tesla MRI scanners according to unified ADNI monitoring protocols*(49)*. Structural MRI and resting-state functional MRI (rsfMRI) scans were acquired during the same visit. High-resolution structural MRI scans were recorded using a T1-weighted MP-RAGE sequence at a voxel size of 1 $mm^3$ and a TR of 2300 ms. For rsfMRI scans, volumes were acquired with under the following parameters: voxel size of 3.3 $mm^3$, TE of 30 ms, TR of 3000 ms, flip angle of 80°, and a minimum of 140 volumes.

*Image processing:*

All data obtained from the ADNI database were converted from DICOM to NIFTI format using the dcm2niix function in the MRIcroGL application*(50)*. Any nonbrain tissue, such as the skull and optic nerves, was removed from these scans using the Brain Extraction Tool in the FSL library*(51)*. The fMRI Expert Analysis Tool from this same FSL library was used to discard the first 4 volumes of these scans to allow for magnetization stabilization and then preform motion correction using MCFLIRT rigid body transformations*(52–54)*.

Functional images were then processed using the CONN toolbox, version 22.a*(55)*. Functional images were co-registered to structural T1-weighted images and spatially normalized to the standard Montreal Neurological Institute 152 (MNI) reference space. Functional data were denoised using the default CONN fMRI denoising pipeline*(56)*. Signals were individually filtered with a 0.008 – 0.09 Hz temporal band-pass filter. No spatial smoothening was applied during any of these steps to avoid effacing small areas of significance*(57)*.

Automated anatomical parcellation was manually verified for each subject. Once brains were registered to neuroanatomical atlases, ROI-to-ROI connectivity was assessed between the RSC seeds and other canonical regions within the default mode network (DMN): the medial prefrontal cortex (mPFC), the left and right lateral parietal cortices (LP L and LP R), and the posterior cingulate cortex (PCC). The bilateral RSC seeds each measured 5 mm in diameter and were centered about MNI coordinates $x=\pm6$, $y=-50$, $z=10$, based on expected local activity maxima for this region*(7, 58)*.

*Functional connectivity analyses*:

Anti-correlated activity between RSC seeds and the other DMN regions was assessed by calculating bivariate Pearson's correlations on the BOLD signals recorded between all pairs of ROIs. The resulting ROI-to-ROI coactivation matrices were then filtered to only consider anti-correlations. For each subject, the total number of anti-correlated functional connections was summed across both the left and right RSC and expressed as a percentage of a fully saturated network.
